# Holistic Motor Control of Zebra Finch Song Syllable Sequences

**DOI:** 10.1101/2025.05.04.652139

**Authors:** Massimo Trusel, Junfeng Zuo, Danyal H. Alam, Ethan S. Marks, Therese M.I. Koch, Jie Cao, Harshida Pancholi, Brenton G. Cooper, Wenhao Zhang, Todd F. Roberts

**Affiliations:** Department of Neuroscience, UT Southwestern Medical Center, Dallas TX, USA; Peking-Tsinghua Center for Life Sciences, Academy for Advanced Interdisciplinary Studies, Peking University, China; Lyda Hill Department of Bioinformatics, UT Southwestern Medical Center, Dallas TX, USA; Department of Psychology, Texas Christian University, Fort Worth TX, USA; Peter O’Donnell Jr. Brain Institute, UT Southwestern Medical Center, Dallas TX, USA

## Abstract

How brain circuits are organized to skillfully produce learned sequences of behaviors is still poorly understood. Here, we functionally examine how the ‘cortical’ song premotor region HVC, which is necessary for zebra finch song, controls the sequential production of learned song syllables. We find that HVC can generate the complete sequence of learned song syllables independently of its main synaptic input pathways. Thalamic input to HVC is permissive for song initiation but it is not required for transitions between syllables or completing song. We show that excitation of HVC neurons during song reliably causes vocalizations to skip back to the beginning of song, reminiscent of a skipping record. This restarting of syllable sequences can be induced at any moment in song and depends on local circuits within HVC. We identify and computationally model a synaptic network including intratelencephalic premotor and corticostriatal neurons within HVC that are essential for completing song syllable sequences. Together, our results show that the learned zebra finch song is controlled by a ‘cortical’ sequence-generating network in HVC that, once started, can sustain production of all song syllables independent of major extrinsic input pathways. Thus, sequential neuronal activity can be organized to fuse well-learned vocal-motor sequences, ultimately achieving holistic control of this naturally learned behavior.

## Main

It can be challenging to draw direct links between naturally learned behaviors and the brain circuits that are required for their expression. The precise control of learned birdsong provides a tractable model in which to draw these links. Songbirds are one of the few groups of animals, aside from humans, that learn their vocalizations through imitation^1,2^. Moreover, production of birdsong is controlled by dedicated pallial and basal ganglia neural circuits^3^.

Zebra finches learn a single courtship song, characterized by the serial and stereotyped production of several complex song syllables, and they engage in extensive daily practice to maintain expert performance of this song. Sparse sequential neuronal activity in the pallial song nucleus HVC likely underlies the production of zebra finch song.^4–8^ However, how neural sequences in HVC contribute to the progression of song syllables is still not well understood. Several lines of evidence support the idea that song control may involve reciprocal loops spanning the brainstem, thalamus and pallium^9–12^ (Extended Data Fig. 1A), while other research suggests that HVC may be capable of generating neural sequences for song production more autonomously^5,6,13,14^.

Research in just the past few years has indeed inspired various models of how adult song is controlled, including I) that sequential activity in HVC can sustain progression through all song syllables independently of instructive afferent inputs^10,13–15^, II) that input pathways link shorter neural sequences at syllable or other vocal parameter boundaries^12,16,17^, or III) that HVC sequences are continuously updated by instructive afferent input^18^. Much of the research in this area has relied on correlations between song and electrophysiological recordings or on non-selective circuit manipulations including electrical stimulation, cooling of brain regions, and electrolytic lesioning.

Here, we combine a series of specific cell-type, circuit, and pathway manipulations with synaptic mapping and computational modeling to causally test and examine how neural sequences contribute to the progression of song syllables. This research reveals that, barring a permissive thalamic input important for song initiation, HVC can independently propagate activity for production of all song syllables and that this network relies on two synaptically interconnected classes of HVC projection neurons.

## Optogenetic restarting of song

Electrical stimulation of HVC during singing is reported to have varied effects on song production, including distortion of syllable acoustic features, truncation of song, and the occasional restarting of song soon after song truncation ^19–21,22^. However, electrical stimulation studies are difficult to interpret because it is not possible to restrict stimulation to specific cell types or to cells within a small spatial volume, the stimulated population of neurons is highly dependent on electrode placement, and there is inevitable antidromic and orthodromic activation of neurons and passing axons^23^.

To circumvent these issues, we selectively controlled HVC neuronal activity using viral expression of the excitatory opsin ChRmine^24^ (n = 6 birds, **Fig. 1A**; **Extended Data Fig. 1B**). Birds were bilaterally implanted with fiber optics over HVC, and syllable detection software was used to perform closed-loop optogenetic manipulations while they were freely singing (see Methods). We found that light stimulation (10-50ms) reliably caused song truncation, seen as a rapid decrease of sound amplitude and disruption in syllable acoustic features (stimulation outcome probability: 86.76±3.61% truncation, 10.23±3.66% pause+continuation; latency to silence from onset of stimulation: 66.55±4.07ms, AVG±SEM in 6 birds, **Fig. 1B-G**; **Extended Data Fig. 1C-G**). Stimulation-evoked truncation was followed by the rapid restarting of the song motif (the stereotyped sequence of syllables is referred to as the “motif” and zebra finches string together several repeats of the motif in a song “bout”, which is usually preceded by production of “introductory notes”, **Fig. 1B-D**; **Extended Data Fig. 1C**). We found that birds often restart their song from the beginning, either with one or two introductory notes followed by the motif or directly back to the first syllable of the motif, and that this resetting behavior occurred with high probability, independent from when in the song the optogenetic stimulation was triggered (**Fig. 1H**, **all post-truncation trials reported in** **Extended Data Fig. 1H**). When computed across all stimulation trials and normalized by the likelihood of birds to chain multiple motifs in series, the probability of a stimulated motif to be immediately followed by another motif was 108.6±4.9%, suggesting that the optogenetic perturbation caused the song to restart from the beginning of the motif without prematurely ending the song bout (**Fig. 1I**, not-normalized motif reset probability computed over all trials 69.16±1.82%; **Extended Data Fig. 1I**). It has been shown that sensory feedback to HVC is gated off during singing^25–27^ and we find that the rapid restarting of the motif is consistent with internal neural circuit dynamics rather than sensory feedback (**Fig. 1J** median = 135.8±25.76 ms, lowest quartile = 87.58±15.52 ms).

**Fig. 1.**
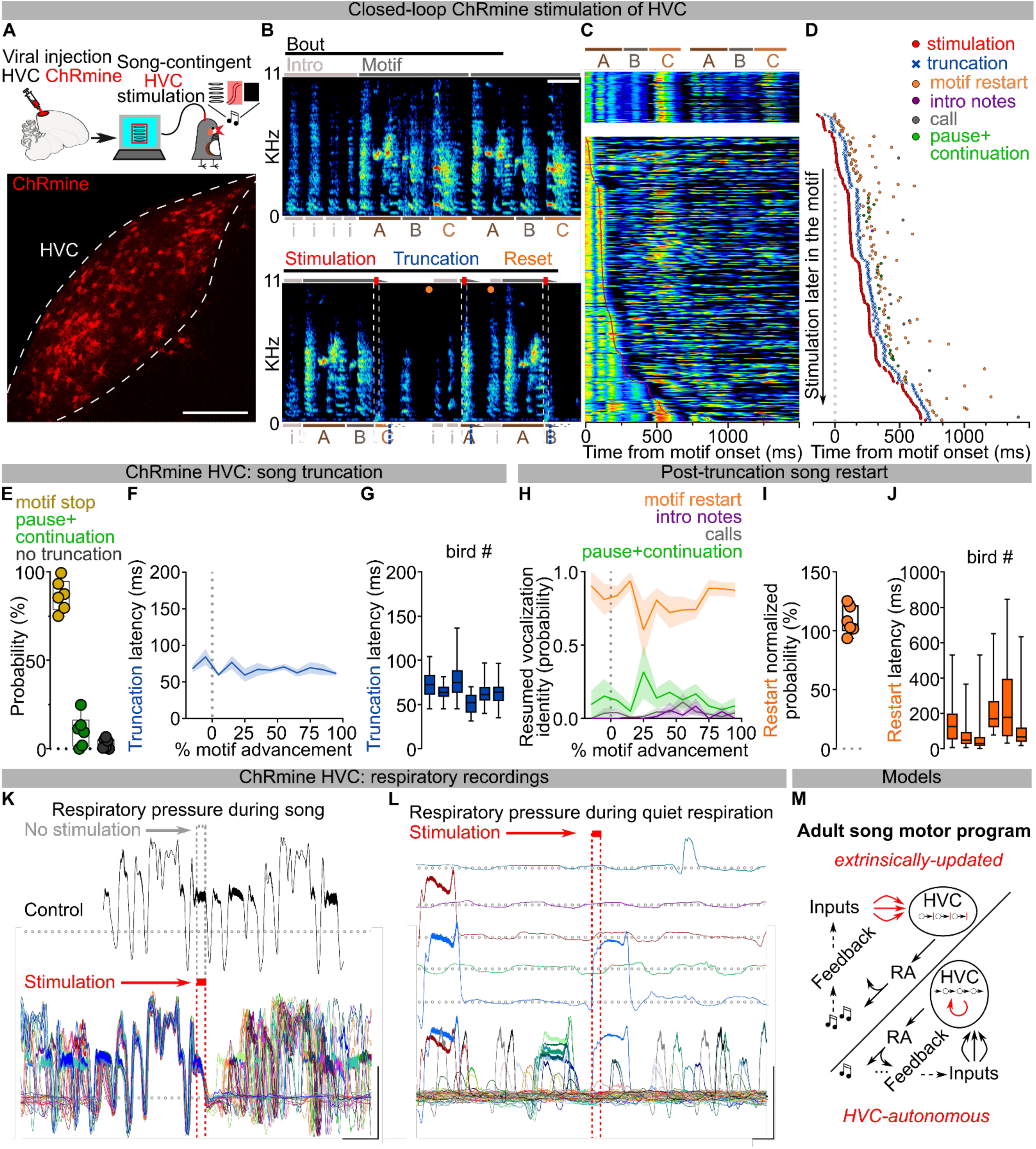
Optogenetic excitation of HVC causes truncation and restarting of the song motif. **A)** (top) Schematic of closed loop song-contingent light stimulation of HVC neurons expressing the excitatory opsin ChRmine; (bottom) sample image of HVC expressing ChRmine (scalebar 200µm). **B)** (left) Spectrograms of normal (top) and stimulated (bottom) song bouts (scalebar 200ms). Horizontal lines identify the bouts’ (black), motifs’ (dark gray) and syllables’ boundaries, introductory notes marked as “i” (light gray) are followed by the syllables (marked with letters A,B,C) composing the motif. Light stimulation (red bars: 10ms light pulses, white dashed lines on the spectrograms mark the segments of song being stimulated) causes motif truncation of syllables (blue dashed lines overlaying letters A-C, residual expected segment representing the missing portion of the truncated syllable is shown as dashed contour; motif truncation represented by the motif line being truncated at an angle). Truncation is followed by motif restart (orange dot). **C)** Stacked song amplitude plot showing amplitude of unstimulated (top) and all stimulated motifs (bottom) from the bird in (B), ordered by timing of stimulation onset (red line, red arrow points at start). **D)** Plot reporting a subset of stimulated motifs’ latency to optogenetic stimulation (red circle), motif truncation (blue “x”), and restart of a motif (orange), intro notes not followed by a motif (purple), calls (grey) or continuation of the motif after a pause (green) normally not present in the unstimulated motif within 1 second following stimulation. **E)** Box and scatter plot reporting the probability of motif stop (okra), pause and continuation of the motif (green) or absence of syntactic perturbation (gray) after the light stimulation (n=6 birds). **F)** Average latency ±SEM to motif truncation in response to the light stimulation (n=6 birds) across the motif (% motif advancement, bins = 10% motif advancement). **G)** Box plot (quartiles and 5-95%) reporting the truncation latencies computed across all stimulations for each bird. **H)** Average ±SEM probability of post-truncation resumption category (motif restart (orange), intro notes not followed by a motif (purple), calls (grey), resumption of the motif after a pause (green)) in response to the light stimulation delivered at different latencies throughout the motif (bins = 10% motif advancement). **I)** Box and scatter plot reporting the normalized probability of post-truncation motif restart for each bird (see methods). **J)** Box plots (quartiles and 5-95%) representing the motif restart latency of each bird. **K)** Subsyringeal pressure recordings (dotted line indicates ambient pressure, deviations above ambient pressure indicate expiration and below indicate inspiration) from a bird subjected to optogenetic stimulation of HVC neurons during song, aligned at the onset of stimulation, displaying the rapid loss of pressure caused by the stimulation (top: control unstimulated trace, aligned to bottom: 34 motif traces aligned at stimulation onset (red dashed bar, 50ms); corresponding gray dashed bar highlights the corresponding point in the unstimulated control motif waveform; scalebar: 200ms, 0.5A.U.). **L)** Subsyringeal pressure recordings from the same bird in panel (K) stimulated during quiet respiration or calls (top: sample traces displaying the lack of effect of stimulation delivered at different moment during the inspiratory-expiratory cycle, or during onset of a call; bottom: 56 traces aligned at the stimulation onset, scalebar: 200ms, 0.5A.U.). **M)** Schematic of the 2 proposed possible scenarios in which song progression is controlled through extrinsic updates to HVC activity or controlled more autonomously by HVC.

To better understand how our circuit manipulations affect motor control of song, we recorded subsyringeal air sac pressure during HVC optogenetic stimulations. First, we found that the effect of optogenetic stimulation in HVC is strongly state dependent. When applied during quiet respiration it did not cause birds to vocalize or affect respiratory patterns. By contrast, stimulation during singing caused rapid cessation of expiration during ongoing syllables and we anecdotally observed that repeated optogenetic activation while birds were singing could time-lock respiratory activity and vocalizations (**Fig. 1K-L**; **Extended Data Fig. 1J**). Second, syllable truncations resulted from significant (>2SDs) respiratory pressure deviations within 36.35±4.00ms of light onset, ∼30ms before vocalizations were acoustically truncated, consistent with previous studies^10^ (**Extended Data Fig. 1K**). Third, we found that at least some of the optogenetic stimulation trials in which birds do not quickly restart singing could be the result of the stimulation inducing apnea. Thus, in some cases optogenetic activation suppresses involuntary respiration, which effectively blocks the re-initiation of song (apnea duration 588.2±216.8ms; **Extended Data Fig. 1L**).

Together, these data indicate that HVC neurons can control downstream steady-state respiration circuitry in a state dependent manner and that, once HVC is engaged, stimulation interrupts the chain of activity in HVC resulting in abrupt song truncation and resetting of the motif back to its initial state. These attributes (state dependence and resetting of patterned movement sequences for song following perturbation) are reminiscent of central pattern generating (CPG) networks described in the invertebrate and vertebrate nervous systems^28,29^. Another defining feature of CPGs is that once initiated they can produce patterned activity in the absence of instructive patterned input. The seemingly automatic and rapid restarting of song hints that extrinsic inputs to HVC may function permissively, rather than instructively, in song production (**Fig. 1M**), thus raising the possibility that HVC produces the neuronal sequences for song in the absence of instructive patterned input and may function as a pattern generating network for song syllable sequences.

## Thalamic Input is Permissive for Song

Thalamic input to HVC from the thalamic Nucleus Uvaeformis (Uva) is one likely source of instructive or permissive signals important for song sequence progression^13,18,30,31^. Electrical stimulation of Uva is reported to cause motif truncation at syllable boundaries^12^, suggesting that Uva’s inputs to HVC are instructive for motor programs to transition from one syllable to the next. To test this idea using pathway selective manipulations we first used optogenetic excitation of Uva’s axon terminals in HVC via viral expression of eGtACR1, an opsin we have found to potently drive excitation of axon terminals in zebra finches^31^ **(Extended Data Fig. 2A)**. Light stimulation of eGtACR1-expressing Uva-HVC terminals drives strong transient increases in HVC activity (**Fig. 2A**). In contrast to thalamic electrical stimulation^12^, optogenetic excitation of Uva terminals during singing did not cause motif truncation and left song syntax and spectral characteristics unaffected (**Fig. 2B-C**; **Extended Data Fig. 2B-C**).

**Fig. 2.**
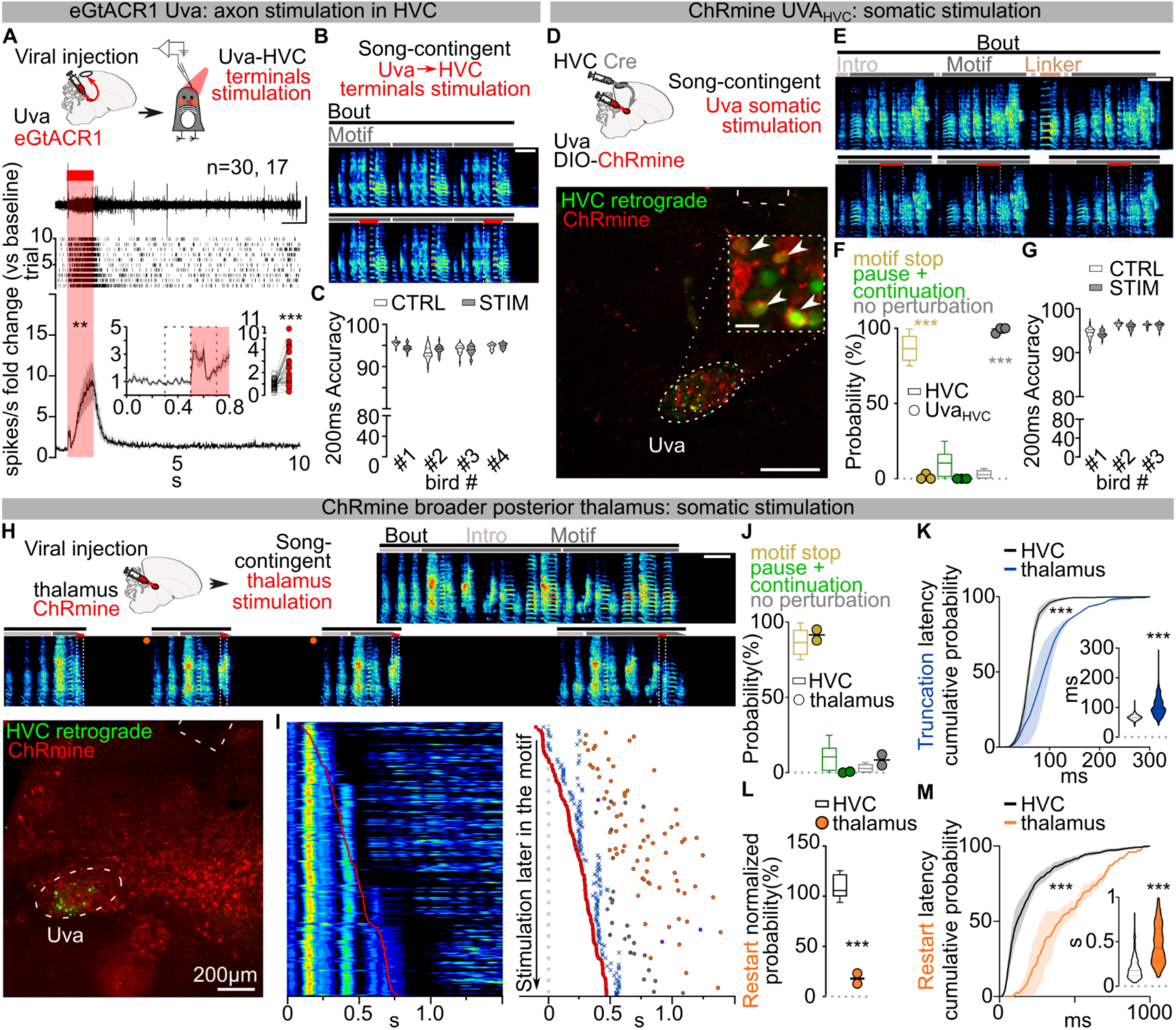
Uva does not instruct transitions between syllables in the song motif. **A)** Schematic of in-vivo recording of HVC multiunit neuronal activity in anesthetized birds expressing eGtACR1 in Uva; sample trace (top), raster plot from 10 stimulation trials (mid) and normalized peri-stimulus time histogram (bottom) reporting the change in multi-unit HVC firing activity in response to light stimulation of eGtACR1-expressing Uva afferents (1s, red bar, two-way ANOVA, F(1,29)=9.096, P=0.0053); inset displays magnified PSTH and scatter plot highlighting and computing the average (per hemisphere) response to the first 200ms light stimulation (red dashed rectangle) compared to the last 200ms baseline (black dashed rectangle, Wilcoxon test, P<0.001; n= hemispheres, birds). **B)** Song-contingent light stimulation of Uva axonal terminals in HVC (optic fiber implanted over HVC): sample spectrogram of unstimulated (top, scalebar 200ms) and stimulated song (bottom, red bars, 200ms ≈10mW bilateral red LED) (spectrogram scale, 0-11KHz, scalebar 200ms). **C)** Violin plots reporting accuracy of song segments with (gray) and without (white) stimulation per each bird (n=4); two-way ANOVA, CTRL vs STIM, F(1,76) = 0.5675, P=0.4536). **D)** (top) Schematic of the experiment using AAV-CRE injection in HVC and AAV-DIO-ChRmine injection in Uva to achieve selective expression of ChRmine in Uva neurons projecting to HVC. (bottom) Image of Uva with ChRmine-mScarlet expression in UvaHVC neurons retrogradely labeled by tracer injection into HVC (scalebar 200µm, inset scalebar 20µm). Dotted white lines at top mark end of implanted fiber optic. **E)** Sample song spectrograms of unstimulated (top, scalebar 200ms) and stimulated (bottom, red bars, 200ms) motifs. **F)** Box and scatter plot reporting the probability of motif stop (okra), pause and continuation of the motif (green) or absence of syntactic perturbation (gray) after the light stimulation (UvaHVC stimulated birds, n=3, filled circles; empty box plots from HVC stimulation in Fig.1E1E reported for comparison; two-way ANOVA F(7,22)=2.549, P=0.0440, Dunnett’s post-hoc, motif stop P<0.001, pause+continuation p=0.4836, no perturbation P<0.001). **G)** Violin plots reporting accuracy of song segments with (gray) and without (white) stimulation for each bird (n=3; two-way ANOVA, CTRL vs STIM, F(1,57) = 3.746, P=0.0579). **H)** Schematic and sample image showing non-selective expression of AAV-ChRmine in Uva and peri-Uva thalamus, followed by thalamic song-contingent stimulation. Sample spectrogram (scalebar 200ms) displaying motif truncation at syllable boundaries caused by thalamic light stimulation (50ms 532nm light pulses, red bars).Sample image illustrating ChRmine expression (red, scalebar 200ms) in Uva (UvaHVC neurons labeled by tracer injection in HVC (green)) and peri-Uva, stimulated by light delivered through the implanted fiber optic (white dashed lines top of image). **I)** (left) plot showing amplitude of all the stimulated (red line) motifs, ordered by time of stimulation in the motif. (right) Plot reporting a subset of stimulated motifs’ latency to optogenetic stimulation (red circle), motif truncation (blue “x”), and restart of a motif (orange), intro notes not followed by a motif (purple), calls (grey) or continuation of the motif after a pause (green) normally not present in the unstimulated motif within 1 second following stimulation. **J)** Box and scatter plot reporting the probability of motif stop (okra), pause and continuation of the motif (green) or absence of syntactic perturbation (gray) after the light stimulation (thalamus stimulated birds, n= 2, filled squares; empty box plots from HVC stimulation in Fig.1E reported for comparison; two-way ANOVA F(7,22)=2.549, P=0.0440, Dunnett’s post-hoc, motif stop P=0.9883, pause+continuation P=0.6761, no perturbation P=0.9728). **K)** Cumulative probability curves reporting the latency to song truncation in response to the light stimulation (average ±SEM of each bird’s curve; thalamus-stimulated birds (blue), HVC-stimulated birds (black) dataset from Fig.1 compared against all experimental groups across the manuscript, 10ms time bins, two-way ANOVA, F(5,17)=4.134, P=0.0122, Dunnett’s post-hoc identifies significant (P<0.05) differences between 60 and 140ms time bins. (inset) Violin plots reporting the latency of motif truncation computed across all the birds (thalamus-stimulated birds (blue), HVC-stimulated birds (white) dataset from Fig.1 compared against all experimental groups across the manuscript; one-way ANOVA, Kruskal Wallis test H(5)=467.6, post-hoc HVC vs. thalamus, P<0.001). **L)** (left) Box and scatter plot reporting the normalized probability of post-truncation motif restart for each bird (thalamus stimulated birds (filled circles); empty box plots from Fig.1I reported for comparison; one-way ANOVA, F(5,16)=9.217, P=0.0003, Dunnett’s post-hoc HVC vs. thalamus: P<0.001). **M)** Same as panel (K) but for latency to motif restart. Two-way ANOVA, F(6,22)=5.850, P=0.0036, Dunnett’s post-hoc identifies significant difference (p<0.05) between 70 and 640ms time bins. (inset) One-Way ANOVA, Kruskal Wallis test H(6)=245.5, post-hoc HVC vs. thalamus P<0.001).

The lack of effects on song, even with prolonged stimulation, is surprising and prompted us to test whether direct optogenetic excitation in Uva disrupts song. We selectively expressed the excitatory opsin ChRmine in Uva neurons projecting to HVC (Uva_HVC_) using an intersectional viral strategy (**Fig. 2D**). We found that even directly stimulating Uva_HVC_ neurons failed to cause song truncation and restarting (1.15±1.15% motif stop, no effect: 98.85±1.15%; **Fig. 2E-F**). Moreover, this manipulation also had no detectable impact on the spectral characteristics of song syllables (**Fig. 2G**; **Extended Data Fig. 2D**).

One possibility is that manipulations like electrical stimulation may drive truncation at syllable boundaries via off-target effects, like recruiting nearby thalamic regions or fibers of passage. Uva is located within the posterior commissure, which connects midbrain regions critical for vocalizations, audition and vision, and is immediately adjacent to the RA fiber tract which transmits descending motor commands for song (**Supplementary video 1)**. Critically, neurons in and surrounding Uva relay visual information to the forebrain^32^ and sudden visual stimulation with a stroboscope elicits orienting responses in zebra finches^33,34^ that result in motif truncations at syllable boundaries, like those reported by Moll et al., 2023.

To distinguish between fibers of passage, or visual relay, we attempted to mimic the effects of electrical stimulation by non-selectively expressing ChRmine in Uva and the surrounding thalamus (**Fig. 2H**). We found that this broader thalamic stimulation resulted in reliable motif truncation at syllable boundaries (91.48±3.62% motif stop, 0.35±0.35% pause+continuation, **Fig. 2I-J**). In contrast to optogenetic stimulation in HVC or the Uva-HVC pathway, optogenetic stimulation of the broader thalamus caused birds to momentarily stop movement and blink during singing and also during non-singing states, suggesting that this manipulation causes a visually evoked orienting response, perhaps mimicking responses to strobe light visual stimulation. Consistent with this, broader thalamic stimulation resulted in significantly longer truncation latencies than direct HVC stimulation (**Fig. 2K**; **Extended Data Fig. 2E-F**) and predominantly resulted in cessation of singing rather than resetting of song (**Extended Data Fig. 2G**). In the few instances when birds returned singing (**Fig. 2L**; **Extended Data Fig. 2H**), the motif reset latency was significantly longer than what we observed when stimulating HVC (**Fig. 2M**; **Extended Data Fig. 2I**). Together, these findings suggest that electrical stimulation-triggered song truncations might be the result of off-target stimulation of the broader peri-Uva thalamus, and that Uva may not play an instructive role in HVC syllable sequence progression.

We next examined if Uva could play a permissive, rather than instructive, role in song production. Electrolytic lesions of Uva or peri-Uva regions can abolish courtship song production^18,30,35^. However, electrolytic lesions non-selectively ablate neurons as well as damage axonal fibers in and around Uva, including axons from nucleus RA in the posterior commissure essential for song^21,36^ (**Supplementary video 1)**. To minimize damaging axonal tracks, we performed bilateral excitotoxic lesions of Uva using a cocktail of ibotenic and quisqualic acid (n = 13 birds). This strategy yielded 3 main outcomes: (i) complete or almost-complete lesions of Uva (99.56±0.44% Uva lesioned) that also included the peri-Uva thalamus that resulted in birds that could no longer sing their motif after the lesions; (ii) large peri-Uva thalamic lesions that mostly spared Uva (10.75±3.75% Uva lesioned) that resulted in birds that also could no longer sing their song motif; and (iii) almost complete Uva lesions (87.50±6.95% Uva lesioned) that spared the broader peri-Uva thalamus and resulted in birds that could sing their motif within ∼1 week following lesion (**Fig. 3A-C**; **Extended Data Fig. 3A**). This last group of birds demonstrates that HVC can drive production of the entire song motif even when Uva is significantly lesioned. Nonetheless, we found that these birds chained significantly fewer motifs together in each song bout (**Fig. 3D**) and they would often fail to produce their song motif after singing introductory notes (**Fig. 3E**). These findings are consistent with Uva lesions disrupting the ability of birds to initiate courtship song performances to female birds^30^, and suggest that Uva may be permissive for the initiation of the song motif, rather than instructing syllable transitions within the motif.

**Fig. 3.**
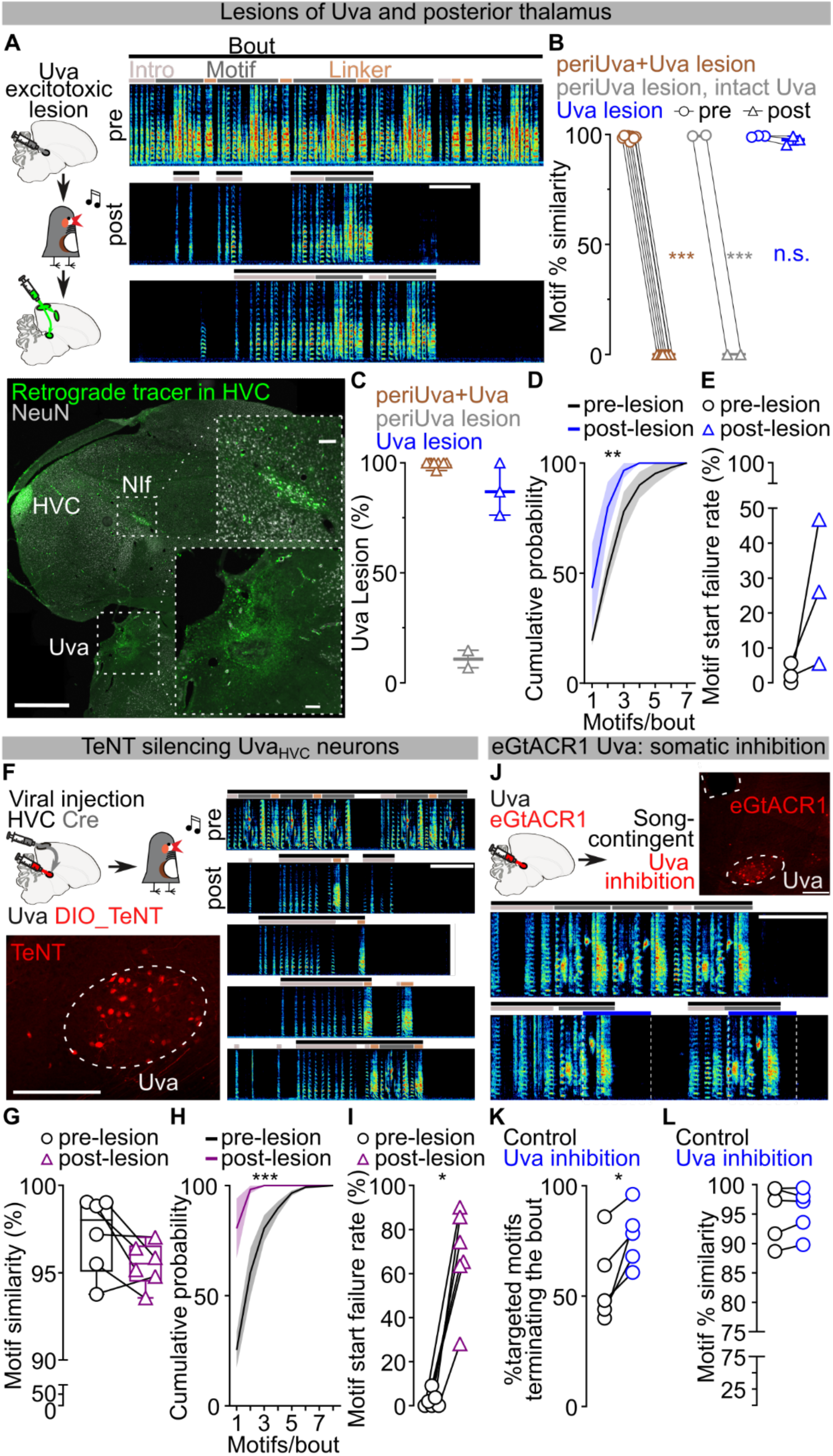
Uva is permissive for motif initiation. **A)** Schematic, sample image (scalebar 1mm, insets 100µm) and spectrograms (scalebar 1s) reporting the effect of excitotoxic lesion of Uva and peri-Uva thalamus. Spectrograms show song pre and post bilateral Uva lesion. NeuN immunofluorescence (gray) and retrograde tracing from HVC (green) highlights surviving neurons in NIf but not in and surrounding Uva. **B)** Motif self-similarity (lack of motif scored as 0) of birds before (circles) and 1-2 weeks after (triangles) the excitotoxic lesion of Uva and peri-Uva thalamic nuclei (peri-thalamus+Uva (brown; n=8, paired t-test, P<0.001); peri-thalamus excluding Uva (gray, n=2, paired t-test, P=0.0013); Uva excluding perithalamic areas (blue, n=3, paired t-test, P=0.2082). **C)** The % lesion of Uva in the 3 experimental groups (peri-thalamus+Uva (brown), peri-thalamus excluding Uva (gray), Uva excluding the larger perithalamic areas (blue)). **D)** Cumulative probability of motifs/bout sung by the birds before (black) and 30d after (blue) with excitotoxic lesion of Uva that spared peri-Uva thalamic regions (n=3 birds, two-way ANOVA, F(1,14)=12.91, P=0.0029). **E)** The rate of motif start failures before (black circles) and after (blue triangles) the excitotoxic Uva lesions (Wilcoxon test, P=0.2500). **F)** Schematic, sample image (scalebar 200µm) and spectrograms (scalebar 1s) reporting effects of TeNT expression in Uva neurons projecting to HVC. **G)** Motif self-similarity between motifs before viral injection (circles) and the last motifs produced before complete cessation of singing in birds with expression of TeNT in UvaHVC neurons for 1-2 weeks (triangles; n=6 birds, paired t-test, P=0.0659). **H)** Cumulative probability of motifs/bout before (black) and 1-2 weeks after expression of TeNT in UvaHVC neurons (purple; last motifs before complete cessation of song; n=6 birds, two-way ANOVA, F(1,40)=43.08, P<0.001). **I)** Rate of motif start failures before (black circles) and after expression of TeNT in UvaHVC neurons (purple triangled, n=6 birds, Wilcoxon test, P=0.0312). **J)** Schematic, sample image (scalebar 200µm), and spectrograms (scalebar 1s) reporting effects of eGtACR1-mediated inhibition of Uva neurons. **K)** Optical inhibition of Uva using eGtACR1 increases song bout terminations (n=5 birds, control (no light), Uva inhibition (light delivered over eGtACR1-expressing Uva neurons for 1 sec; paired t-test, P=0.0238). **L)** Motif self-similarity during optical inhibition of Uva (paired t-test, P=0.5902).

To test Uva’s role in song initiation, we first blocked glutamate release from Uva_HVC_ neurons using viral expression of tetanus neurotoxin (TeNT) (**Fig. 3F**; **Extended Data Fig. 3B**). We found that these birds had progressive difficulty initiating their song on a timeline consistent with viral expression (∼10 - 14 days). TeNT-expressing birds had increasing failures in motif initiation following singing of introductory notes and decreased number of motifs per song bout. Nonetheless, in instances when the motif was initiated, birds consistently produced all song syllables in the motif with high accuracy (**Fig. 3G-I**; **Extended Data Fig. 3C**). This data supports the idea that the Uva-HVC pathway is permissive for initiating learned song motifs rather than instructing song syllable transitions.

To more definitively test this idea, we expressed eGtACR1 in Uva and implanted a fiber optic over Uva to optogenetically silence Uva neurons during singing (**Fig. 3J**; **Extended Data Fig. 2A**). We found that silencing Uva during an ongoing song motif did not disrupt the completion of that motif (**Fig. 3L**), but it reduced the probability of initiating and concatenating a subsequent motif (**Fig. 3L**). In contrast, placing fiber optics over HVC to excite Uva axon terminals expressing eGtACR1 across motif transitions did not suppress initiation of a subsequent song motif (**Extended Data Fig. 3D,E**). Thus if Uva input to HVC is excited birds can continue the ongoing motif and string other motifs together. If instead it is inhibited birds still complete the ongoing motif, yet have difficulty starting the next song motif. Together these findings indicate that the Uva-HVC pathway is critical for initiating song motifs but not needed for birds to string together syllables within the song motif.

## Pallial afferents are not necessary for song

In addition to thalamic input, HVC receives excitatory input from three auditory and premotor pallial regions that play important roles in song learning: Nucleus Interfacialis (Nif), nucleus Avalanche (Av), and medial magnocellular nucleus of the anterior nidopallium (mMAN). We examined the role of each of these afferent pathways on adult song performance. Stimulation of eGtACR1-expressing axon terminals in HVC from any of these regions significantly increased HVC multiunit firing activity (**Extended Data Fig. 4A,G,M**). However, 200ms or 1s-long song-contingent light stimulation of any of these input pathways failed to affect spectrotemporal motif characteristics, like what we observed for stimulation of Uva terminals (**Extended Data Fig. 4A-R**). We therefore tested whether any of these afferents are necessary for adult song performance. To avoid compensation of remaining input pathways, we consecutively and bilaterally lesioned mMAN, NIf, and Av in the same birds. We coupled these lesions with lesion of lMAN, the main output of the song system’s cortico-basal ganglia-thalamo- cortical circuit. Bilateral lesions of these nuclei (mMAN: 100.00±0.00%; NIf: 92.90±4.03%, Av: 100.00±0.00%, lMAN: 82.50±7.67%, **Extended Data Fig. 5A-I**) caused only a temporary decrease in motif quality. The song motif quickly recovered to its pre-lesioned state (**Extended Data Fig. 5J)** and, unlike Uva lesions, these lesions did not impact the number of motifs per bout or cause disruptions in motif initiation (**Extended Data Fig. 5K,L**). These results demonstrate that HVC can generate the sequential activity necessary for completing song independent of its main excitatory synaptic afferents from Nif, Av, and mMAN. These results provide further support for the idea that HVC circuit activity autonomously generates the premotor pattern necessary for song production.

## Song pattern generating network is limited to HVC

We next sought to further define the circuit boundaries of this song pattern-generating network by asking whether HVC’s downstream target regions are critical to pattern generation. We reasoned that the temporal kinetics of restarting song after truncation provides a sensitive behavioral read-out of the pattern generating network. Comparing motif reset latency and probability upon optogenetic perturbation of HVC or its output areas could therefore clarify whether those neural circuits are involved in song pattern generation or simply relay the patterned output. We expect stimulation of relay nodes outside the pattern generating network result in song interruptions with high probability and low latency, but lead to motif reset with lower probability and longer latency. Disruption of the pattern generating network should produce truncation and reset latencies similar or faster than those observed upon broad optogenetic stimulation of HVC. HVC has two major output pathways, the descending song motor pathway through the pallial song region RA and the palliostriatal pathway through Area X, emerging from HVC_RA_ and HVC_X_ neurons, respectively.

We first bilaterally expressed ChRmine in either Area X or RA and light stimulated each region in freely singing birds. Driving Area X neurons very rarely caused motif truncation (truncation probability: 2.85±1.59%, no effect: 97.15±1.59%, **Fig. 4A,B**; **Extended Data Fig. 6A**). The few truncations we observed occurred at syllable boundaries and were significantly delayed (latency: 146.70±36.66ms) compared to the uniform song truncations observed with HVC optogenetic stimulation. Nonetheless, stimulation of Area X neurons did consistently cause a modest increase in the entropy (noisiness) of stimulated syllables (**Fig. 4C,D**; **Extended Data Fig. 6B,C**), consistent with the basal ganglia pathway injecting variability into the song motor program.

**Fig. 4.**
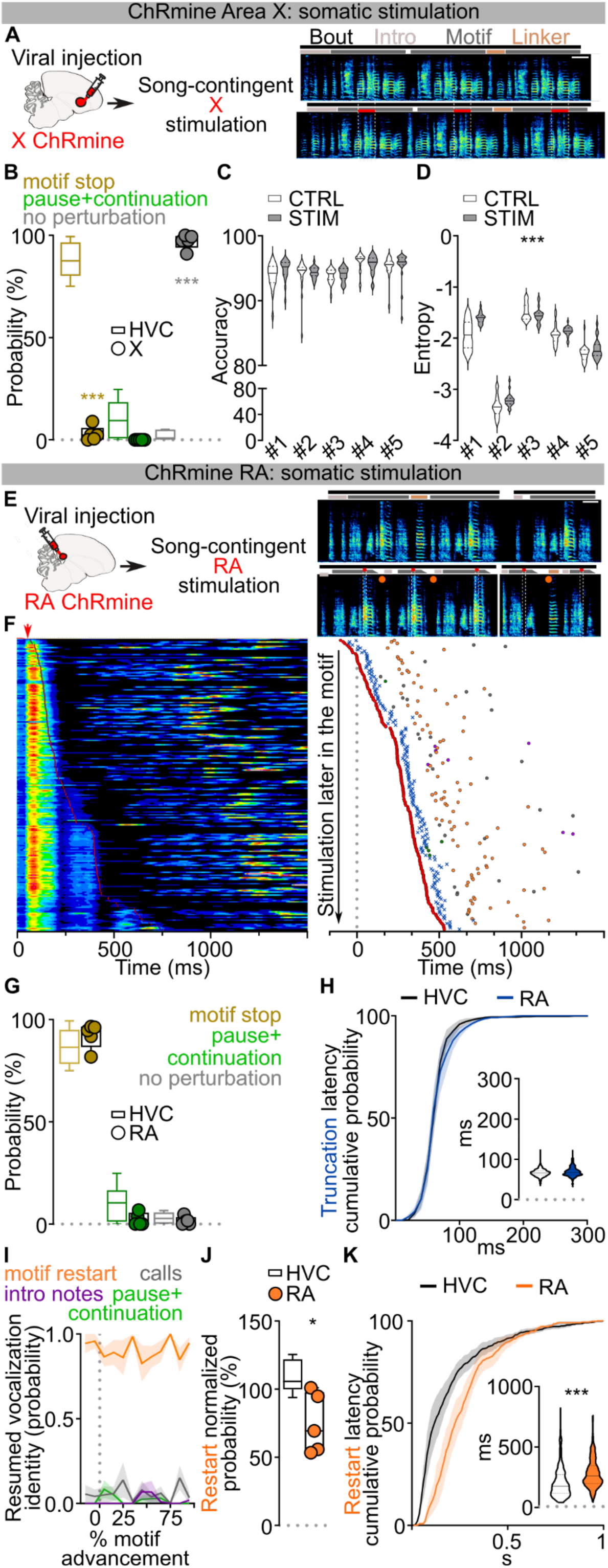
Pattern generation circuit confined to HVC. **A)** Experiment design and song spectrogram showing song-contingent light stimulation of Area X neurons expressing the excitatory opsin ChRmine (top: control, scalebar 200ms; bottom: stimulated, 200ms 532nm light pulses, red bars). **B)** Probability of motif truncation (okra), pause and continuation of the motif (green) or absence of syntactic perturbation (gray) after the light stimulation of Area X (X stimulated birds, n=5, filled circles; empty box plots from HVC stimulation in Fig.1E reported for comparison; two-way ANOVA F(7,22)=2.549, P=0.0440, Dunnett’s post-hoc, motif stop P<0.001, pause+continuation P=0.3117, no perturbation P<0.001). **C-D)** Violin plots reporting accuracy and entropy of song segments with (gray) and without (white) stimulation for each bird (accuracy: n=5, two-way ANOVA, CTRL vs STIM, F(1,95)=1.150, P=0.2862; entropy: n=5, two-way ANOVA, CTRL vs STIM, F(1,95)=16.63, P<0.001). **E)** (left) Schematic of the song- contingent light stimulation of RA neurons expressing the excitatory opsin ChRmine, and (right) spectrograms representing normal (top, scalebar 200ms) and stimulated song (bottom, 10ms 532nm light pulses, red bars) displaying light-evoked motif truncation of syllables followed by motif restart. **F)** (left) stacked song amplitude plots showing all the stimulated motifs, ordered by the timing of stimulation from the motif onset (red line). (right) Plot reporting a subset of stimulated motifs’ latency to optogenetic stimulation (red circle), motif truncation (blue “x”), and restart of a motif (orange), intro notes not followed by a motif (purple), calls (grey) or continuation of the motif after a pause (green) normally not present in the unstimulated motif within 1 second following stimulation. **G)** The probability of motif stop (okra), pause and continuation of the motif (green) or absence of syntactic perturbation (gray) after the light stimulation (RA stimulated birds, n=5, filled circles; empty box plots from HVC stimulation in Fig.1E reported for comparison; two-way ANOVA F(7,22)=2.549, P=0.0440, Dunnett’s post-hoc, motif stop P=0.8821, pause+continuation P=0.4901, no perturbation P=0.9944). **H)** Cumulative probability curves reporting the latency to song truncation in response to the light stimulation (average ±SEM of each bird’s curve, blue: RA-stimulated birds, black: HVC-stimulated birds dataset from Fig.1 compared against all experimental groups across the manuscript, 10ms time bins, two-way ANOVA, F(5,17)=4.134, P=0.0122, Dunnett’s post-hoc P>0.05). (inset) latency of motif truncation computed across all the birds (blue: RA-stimulated birds, white: HVC-stimulated birds dataset from Fig.1 compared against all experimental groups across the manuscript; one-way ANOVA, Kruskal Wallis test H(5)=467.6, post-hoc HVC vs. RA P>0.9999). **I)** Average ±SEM probability of post-truncation behavior in response to the RA light stimulation delivered at different latencies throughout the progression of the motif (motif restart (orange), intro notes not followed by a motif (purple), calls (grey), resumption of the motif after a pause (green); bins = 10% motif advancement). **J)** (left) Normalized probability of post-truncation motif restart for each bird (RA stimulated birds: filled circles; empty box plots from Fig.1I reported for comparison; one-way ANOVA, F(5,16)=9.217, P=0.0003, Dunnett’s post-hoc HVC vs. RA: P=0.0245). **K)** Cumulative probability curves reporting the latency to post-truncation motif restart (average ±SEM of each bird’s curve, orange: RA, black: HVC dataset from Fig.1 compared against all experimental groups across the manuscript, 10ms time bins, two-way ANOVA, F(6,22)=5.850, P=0.0009, Dunnett’s post-hoc identifies significant difference (p<0.05) between 60 and 280ms time bins). (inset) Latency to motif restart computed across all the birds (orange: RA stimulated birds, white: HVC birds dataset from Fig.1 compared against all experimental groups across the manuscript; one-way ANOVA, Kruskal Wallis test H(6)=245.5, post-hoc HVC vs. RA P<0.001).

In contrast to Area X stimulation, optogenetic stimulation in RA caused rapid motif truncations with high reliability (92.22±2.69% motif stop, 1.52±1.84% pause+continuation, **Fig. 4E-G**; **Extended Data Fig. 6D**). These truncations exhibited uniform latency across song, similar to stimulation in HVC (**Fig. 4H**; **Extended Data Fig. 6E,F**). Because RA is downstream of HVC in the descending song motor pathway, if it is a core component of the pattern generation network we might expect birds to restart their song as fast, or faster, than when stimulating in HVC, and with equal probability. However, we find that RA stimulation is less likely to be followed with restarting of the motif, and that when it does it takes significantly longer than following HVC stimulation (**Fig. 4I-K**; **Extended Data Fig. 6G-I**). These findings argue that the song pattern generating network is localized to HVC and that RA functions downstream of this network to relay motor commands for song.

To more explicitly test this prediction, we moved upstream by one synapse and examined whether optogenetic stimulation of HVC_RA_ neurons alone would produce the truncation and song restarts with the same timing and reliability as our pan-HVC optogenetic manipulations in Fig. 1. We used an intersectional viral strategy to achieve ChRmine expression only in HVC_RA_ neurons (**Fig. 5A**; **Extended Data Fig. 7A,B**). As anticipated, song-contingent HVC_RA_ stimulation reliably caused rapid truncations throughout song (86.18±5.64% motif stop, 2.46±1.81% pause+continuation, **Fig. 5B,C**; **Extended Data Fig. 7C**). Unexpectedly, the latency to truncation was significantly longer than what we observed with pan-HVC stimulation (**Fig. 5D**; **Extended Data Fig. 7D**). While post-truncation motif reset probability is comparable to that observed with pan-HVC stimulation (**Fig. 5E,F**; **Extended Data Fig. 7E,F**), restart latency was intermediate to the timing of pan-HVC and RA-stimulation (**Fig. 5G**; **Extended Data Fig. 7G-I**). This intermediate timing for song restarting is somewhat surprising because HVC_RA_ neurons are widely hypothesized to be the core network controlling song progression. This prompted us to investigate whether the other main class of HVC projection neurons, HVC_X_ neurons, may contribute to the rapid restart of the song motif.

**Fig. 5.**
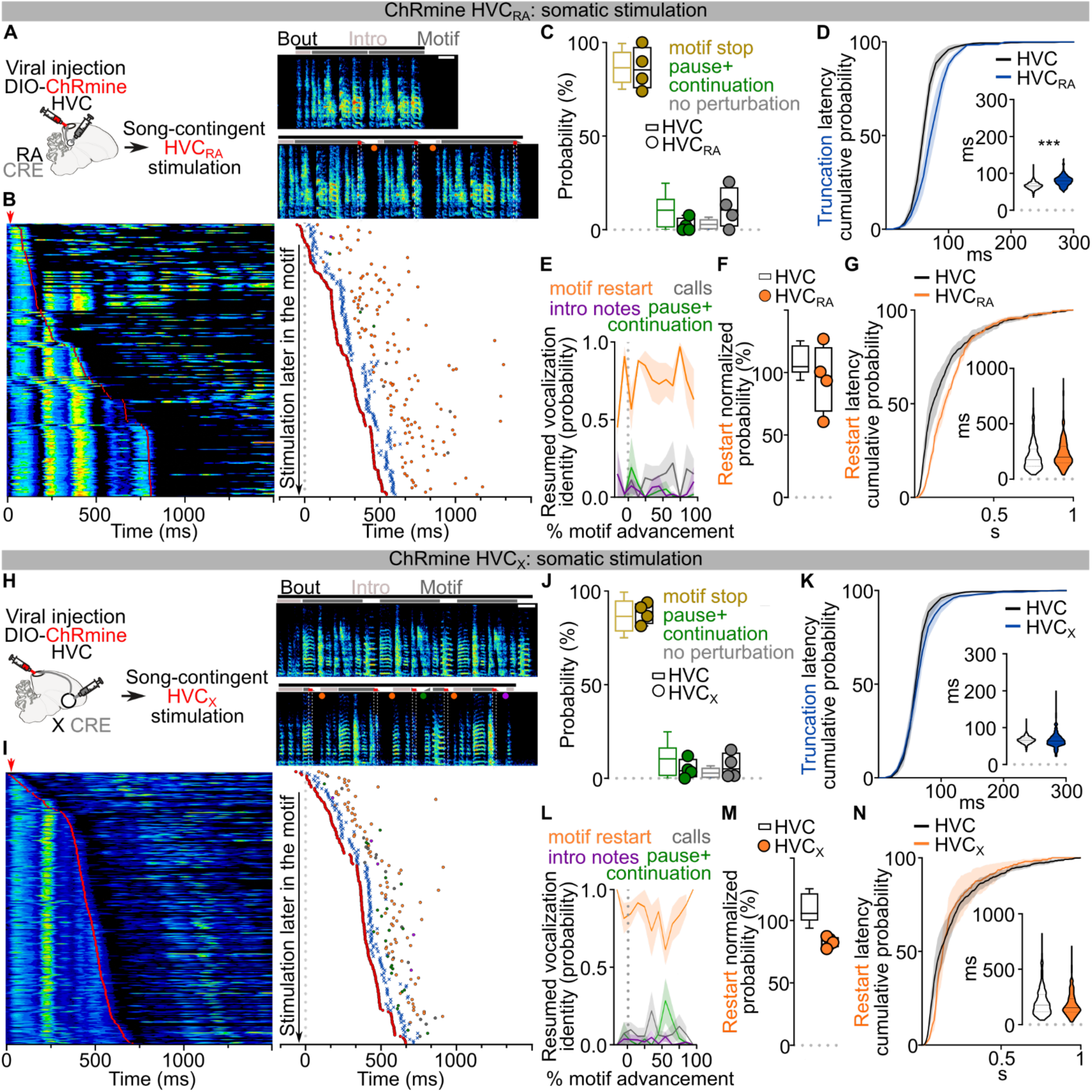
Selective optogenetic stimulation HVC_RA_ or HVC_X_ neurons restarts the song motif. **A)** (left) Schematic of the experiment showing song-contingent in-vivo light stimulation of HVC_RA_ neurons expressing the excitatory opsin ChRmine, and (right) spectrograms representing normal (top, scalebar 200ms) and stimulated song (bottom, 50ms 532nm light pulses, red bars) displaying light-evoked motif truncation of syllables followed by rapid motif restarting. **B)** (left) Stacked song amplitude plot showing all the stimulated motifs, ordered by the timing of stimulation from the motif onset (red line). (right) Plot reporting a subset of stimulated motifs’ latency to optogenetic stimulation (red circle), motif truncation (blue “x”), and restart of a motif (orange), intro notes not followed by a motif (purple), calls (grey) or continuation of the motif after a pause (green) normally not present in the unstimulated motif within 1 second following stimulation. **C)** The probability of motif stop (okra), pause and continuation of the motif (green) or absence of syntactic perturbation (gray) after the light stimulation (HVC_RA_ stimulated birds, n=3, filled circles; empty box plots from HVC stimulation in Fig.1E reported for comparison; two-way ANOVA F(7,22)=2.549, P=0.0440, Dunnett’s post-hoc, motif stop P>0.9999, pause+continuation P=0.5665, no perturbation P=0.7241). **D)** Cumulative probability curves reporting the latency to song truncation in response to the light stimulation (average ±SEM of each bird’s curve, blue: HVC_RA_-stimulated birds, black: HVC-stimulated birds dataset from Fig.1 compared against all experimental groups across the manuscript, 10ms time bins, two-way ANOVA, F(5,17)=4.134, P=0.0122, Dunnett’s post-hoc identifies significant difference (p<0.05) between 60 and 90ms time bins). (inset) Latency of motif truncation computed across all the birds (blue: HVC_RA_-stimulated birds, white: HVC-stimulated birds dataset from Fig.1 compared against all experimental groups across the manuscript; one-way ANOVA, Kruskal Wallis test H(5)=467.6, post-hoc HVC vs. HVC_RA_ P<0.001). **E)** Average ±SEM probability of post-truncation behavior in response to the HVC_RA_ light stimulation delivered at different latencies throughout the progression of the motif (motif restart (orange), intro notes not followed by a motif (purple), calls (grey), resumption of the motif after a pause (green); bins = 10% motif advancement). **F)** Normalized probability of post-truncation motif restart for each bird (HVC_RA_ stimulated birds: filled circles; empty box plots from fig.1I reported for comparison; one-way ANOVA, F(5,16)=9.217, P=0.0003, Dunnett’s post-hoc HVC vs. HVC_RA_: P=0.8071). **G)** Cumulative probability curves reporting the latency to post-truncation motif restart (average±SEM of each bird’s curve, orange: HVC_RA_, black: HVC dataset from Fig.1 compared against all experimental groups across the manuscript, 10ms time bins, two-way ANOVA, F(6,22)=5.850, P=0.0009, Dunnett’s post-hoc identifies significant difference (p<0.05) between 70 and 130ms timebins). (inset) Latency to motif restart computed across all the birds (orange: HVC_RA_ stimulated birds, white: HVC birds dataset from Fig.1 compared against all experimental groups across the manuscript; one-way ANOVA, Kruskal Wallis test H(6)=245.5, post-hoc HVC vs. HVC_RA_ P=0.1157). **H-N)** same as A-G but for HVC_X_ neurons. J) HVC_X_ stimulated birds, n=4, filled circles; empty box plots from HVC stimulation in Fig.1E reported for comparison; two-way ANOVA F(7,22)=2.549, P=0.0440, Dunnett’s post-hoc, motif stop P>0.9999, pause+continuation P=0.9328, no perturbation P=0.9856). K) Average ±SEM of each bird’s curve, blue: HVC_RA_-stimulated birds, black: HVC-stimulated birds dataset from Fig.1 compared against all experimental groups across the manuscript, 10ms time bins, two-way ANOVA, F(5,17)=4.134, P=0.0122, Dunnett’s post-hoc identifies significant difference (p<0.05) at the 70ms time bin; inset, blue: HVC_RA_-stimulated birds, white: HVC-stimulated birds dataset from Fig.1 compared against all experimental groups across the manuscript; one-way ANOVA, Kruskal Wallis test H(5)=467.6, post-hoc HVC vs. HVC_X_ P>0.9999). M) HVC_RA_ stimulated birds: filled circles; empty box plots from Fig.1I reported for comparison; one-way ANOVA, F(5,16)=9.217, P=0.0003, Dunnett’s post-hoc HVC vs. HVC_RA_: P=0.3210. N) Average ±SEM of each bird’s curve, orange: HVC_X_, black: HVC dataset from Fig.1 compared against all experimental groups across the manuscript, 10ms time bins, two-way ANOVA, F(6,22)=5.850, P=0.0009, Dunnett’s post-hoc identifies significant difference (p<0.05) at the 80-90ms time bins; (inset) orange: HVC_RA_ stimulated birds, white: HVC birds dataset from Fig.1 compared against all experimental groups across the manuscript; one-way ANOVA, Kruskal Wallis test H(6)=245.5, post-hoc HVC vs. HVC_X_ P>0.9999.

## HVC_X_ neurons are part of the song pattern generating network

Like HVC_RA_ neurons, HVC_X_ neurons exhibit temporally precise and sparse activity during song^7,27,37^. The role of these neurons in song generation is not known but they have been hypothesized to relay timing activity to the basal ganglia rather than directly contributing to song pattern generation^38–40^. We selectively expressed ChRmine in HVC_X_ neurons (**Extended Data Fig. 7J,K**) and found that stimulation of these neurons during singing reliably triggered song truncations (88.43±2.95% motif stop, 5.02±2.52% pause+continuation, **Fig. 5H-J**; **Extended Data Fig. 7L**). The latency of this truncation matched truncations elicited by pan-HVC stimulation (**Fig. 5K**; **Extended Data Fig. 7M**). Song truncations were followed by rapid song restarts with probability and latency comparable to pan-HVC stimulation (**Fig. 5L-N**; **Extended Data Fig. 7N-P**). Song restart following HVC_X_ stimulation was faster than that from optogenetic HVC_RA_ stimulation (**Extended Data Fig. 7Q,R**). These data indicate that HVC_X_ neurons may be part of the core song pattern generating network, rather than only relaying timing signals to the basal ganglia.

Given that direct optogenetic stimulation of Area X does not result in song truncations (**Fig. 4A-B**), we suspected that perturbation of HVC_X_ neurons synaptic transmission within HVC is the source of truncation and rapid restarting of song. We reasoned that optogenetically stimulating HVC_X_ axon terminals in Area X should cause antidromic excitation of HVC_X_ neurons, thereby recruiting indirect and delayed excitation of axonal collaterals within HVC^19^. If intra-HVC dynamics underlie motif reset, this stimulation should cause delayed song truncation due to the antidromic propagation, yet the motif restart kinetics should be similar to those upon direct HVC stimulation (**Extended Data Fig. 8A**). We first confirmed that optogenetic stimulation of terminals reaching X produces a significant phasic increase in spiking activity in HVC (**Extended Data Fig. 8B).** As anticipated, stimulation of axon collaterals in Area X during singing resulted in delayed motif truncation, albeit with lower probability compared to pan-HVC or HVC_X_ direct stimulation, consistent with the limitations of antidromic propagation^19^ (**Extended Data Fig. 8C-E**). Remarkably however, we found that post-truncation restarting of the song motif had the same probability and latency as stimulating pan-HVC or HVC_X_ neuronal somata (**Extended Data Fig. 8F-H**). This supports the idea that HVC_X_ neurons local connections within HVC can drive song truncations, and dynamics within HVC upon truncation lead to the rapid restarting of song neural sequences.

Previous studies have identified the main synaptic connectivity motif in HVC to be disynaptic reciprocal inhibition between HVC_RA_ and HVC_X_ via local interneurons^41,42^. It is therefore possible that optogenetic stimulation of HVC_X_ neurons would cause a surge of inhibition onto HVC_RA_ neurons, however this doesn’t explain the more rapid truncation and reset of the motif upon HVC_X_ stimulation. To improve our understanding of the synaptic basis for how HVC_X_ neurons may contribute to the song sequence propagation, we mapped the local connectivity of HVC_X_ neurons using opsin-assisted synaptic circuit mapping. Visually targeted whole-cell voltage clamp recordings from HVC_RA_ and HVC_X_ neurons were conducted while optogenetically exciting HVC_X_ axon terminals. Stimulating neurotransmitter release from HVC_X_ neurons evoked excitatory and inhibitory postsynaptic currents in HVC_X_ and HVC_RA_ neurons (**Fig. 6A**; **Extended Data Fig. 9A,B**). We next isolated monosynaptic connections using bath application of tetrodotoxin (TTX) followed by application of 4-aminopyridine (4-AP)^43^. This revealed that HVC_X_ neurons rarely make monosynaptic connections with other HVC_X_ neurons. Instead, they make monosynaptic connections with HVC_RA_ neurons with high probability (**Fig. 6A**). Previous studies using paired recordings suggest that HVC_RA_ neurons also only have sparse connectivity with other HVC_RA_ neurons, yet they are more reliably synaptically connected with HVC_X_ neurons^41^. Altogether, these data support a model in which the two HVC projection neuron classes (HVCPN) form a heterosynaptic network, along with local interneurons, that can holistically sustain song pattern generation.

**Fig. 6.**
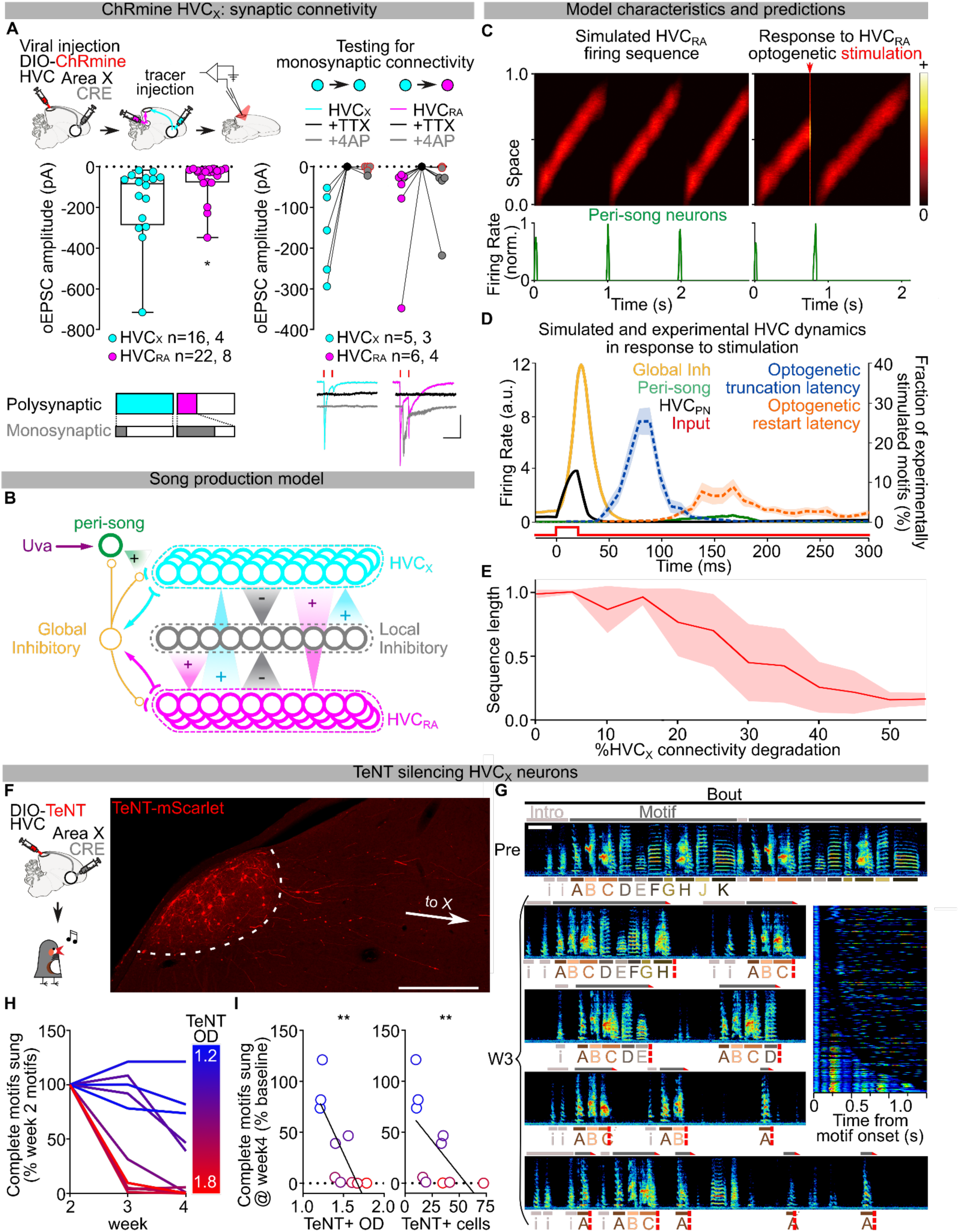
HVC_X_ neurons are part of song pattern generating network and model of HVC. **A)** (top) Schematic for conditional expression of ChRmine in HVC_X_ neurons and retrograde labeling of both HVCPNs (retrogradely labeled) followed by ex-vivo whole cell patch clamp recording; (mid) box and scatter plots showing amplitude of polysynaptic (left) and monosynaptic (right) oEPSCs in HVC_X_ (magenta) and HVC_RA_ (cyan) neurons; for monosynaptic transmission testing: oEPSCs amplitudes after bath application of TTX (black) and 4AP (grey, red outline indicates absence of post-4AP oEPSC, see methods (n= cells, animals); (bottom) sample traces: red lines indicate light stimuli, 1ms; scalebar, 100ms, 100pA; polysynaptic current amplitude, HVC_X_ vs. HVC_RA_, Kruskal-Wallis, P=0.0156); bar charts representing the likelihood of observing polysynaptic oEPSCs in HVC_X_ (cyan) and HVC_RA_ (magenta), and likelihood of a subset of the corresponding oEPSCs to be monosynaptic (gray). **B)** Schematic of model for HVC motif pattern generating circuit. Neurons are depicted by circles. Structured local synaptic projections are shown with cones, unstructured global projections are shown with lines, lines ending with arrows indicate excitatory synapses and lines with circles indicate inhibitory synaptic connections. **C)** Simulated HVC_RA_ sequence for song (left, 3 motifs) and response to simulated optogenetic stimulation truncating a motif and resulting in restart (right, red arrow indicates the stimulation) (bottom) Peri-song neurons activate to restart each motif.. **D)** Modeled dynamics of neuronal activity of HVCPN (black), global interneurons (yellow), and peri-song HVC_RA_ neurons (dark green) in response to excitatory perturbation (red square pulse), aligned and overlayed with experimental AVG±SEM data relative to truncation (blue dashed) and restart (orange dashed) latency upon HVCpan stimulation (dataset from Fig. 1). Experimental data is aligned to onset of optogenetic stimulation. **E)** Mean sequence lengths predicted decrease with the proportion of simulated decreased excitation from HVC_X_ neurons. **F)** Schematic of the experiment and sample image showing conditional expression of TeNT selectively in HVC_X_ neurons (scalebar 500µm). **G)** Spectrograms (scalebar 200ms) representing the progressive shortening and degradation of the motif during week 3 after the viral injection. The inset reports all spectrograms from one day in week 3, aligned at syllable A and ordered by motif length. **H)** Time course reporting the change in number of complete motifs sung by each bird averaged across each week since injection (n motif/day, birds color coded as per heat map on the right reporting the expression amount as normalized optical density). **I)** Scatter plot correlating the relative change in number of motifs/day sung at week 4 as a function of the average optical density of TeNT expression (left, Spearman r=-0.8545, r^2^=0.6127, P=0.0029) or number of TeNT+ somata/50µm thick slice (right, Spearman r=-0.8061, r^2^=0.3533, P=0.0072) or in HVC.

## Model of the HVC pattern generating network

Our findings implicate heterosynaptic transmission between HVC_X_ and HVC_RA_ as a fundamental component of HVC neuronal chain propagation. To test if a circuit organization consistent with this could sustain song progression and restarting, we modeled the network by arranging HVC_RA_, HVC_X_, and local inhibitory neurons uniformly on a “chain” (**Fig. 6B**; **Extended Data Fig. 9C**), ignoring the sparse homosynaptic connections within either class of excitatory projection neuron^42^. Synaptic weights in our model are symmetric and the weights of excitatory and inhibitory connections decay with the distance along the chain. A pool of inhibitory neurons is driven by excitatory neurons and provides global inhibitory feedback to excitatory neurons. We have recently shown that Uva makes prominent monosynaptic connections with HVC_RA_ neurons^31^. Therefore, in this model, song initiation is mediated by excitatory input from Uva onto a recently identified functional class of “peri-song” HVC_RA_ neurons which become active just before song onset and are inactive during singing^44^ (**Fig. 6B**).

We find that setting a sufficiently strong local inhibitory tone allows stable propagation of activity along the chain. Local inhibitory activity spatially lags excitatory activity in the direction of sequence propagation, thus providing stronger inhibition to excitatory neurons at earlier positions in the chain, effectively pushing excitatory activity forward along the chain (**Fig. 6C**; **Extended Data Fig. 9D**, **Supplementary video 2,3**). Weakened excitatory synapses at the end of the chain stop sequence propagation and release peri-song neurons from inhibition, leading to spontaneous restarting of the song motif if excitatory drive from Uva is intact (**Fig. 6B,C**). We then tested how this circuit architecture would respond to strong synchronous excitation, mimicking our optogenetic manipulations (**Fig. 6C**). One possibility is that the activation function of global inhibitory neurons is steeper than excitatory neurons, and the excitatory and global inhibitory neurons are strongly connected in a parameter regime called inhibition-stabilization^45^. Thus, synchronous excitatory activity causes widespread and strong feedback from inhibitory neurons that blocks sequence propagation. Once the sequence is truncated, it can be spontaneously restarted via peri-song neurons, like restarting the song after its natural ending (**Fig. 6C,D**; **Extended Data Fig. 9E**; **Supplementary video 2,3**). Unexpectedly, we found that the parameter settings that allow the model to generate a moving neural sequence, optogenetic truncation, and spontaneous restarting of sequence generation also resulted in timing that qualitatively matched our experimental results. The timing for song truncation and spontaneous restarting from the model matched what we observed following optogenetic manipulations of the HVC network (**Fig. 6D**). Therefore, modelling demonstrates that an inhibition-stabilized pattern generating circuit captures our behavioral results following circuit perturbations, and an emergent property of the model is that it matches the behavioral timing of song truncation and restart.

Lastly, we used this model to further examine the role of HVC_X_ neurons for motif sequence propagation. Although previous studies have suggested that HVC_X_ neuronal lesion leaves song intact^39,40^, our model simulations predict that if we stochastically weaken HVC_X_ neurons contribution to the chain, it will result in premature truncating of neural sequence propagation, and spontaneous restarting of the motif (**Fig. 6B,E,F**, **Supplementary video 2,3**). To test these predictions, we suppressed excitatory synaptic transmission from HVC_X_ neurons using selective viral expression of TeNT (**Fig. 6F**). In contrast to the effect of expressing TeNT in Uva, which resulted in birds producing their song motif in an all or none fashion (**Fig. 3F-I**), we found that TeNT in HVC_X_ neurons caused birds to progressively increase the likelihood of prematurely truncating their song motifs (**Fig. 6F-I**, **Extended Data Fig. 9G-I**). Birds exhibited song truncations both within and between syllables (**Fig. 6G**; **Extended Data Fig. 9J**). The truncations occurred progressively earlier in the motifs, with timelines consistent with viral expression dynamics, and with effects directly proportional to the amount of TeNT expression (**Fig. 6I**; **Extended Data Fig. 9G,K**). Our model also predicts that, if the “song behavioral state” is active, the song motif should restart with a time course matching what was observed following optogenetic excitation of HVC_X_ neurons. We found that birds would frequently restart their songs following premature motif truncations and that the latency of these restarts matched those observed upon direct optogenetic excitation of HVC_X_ neurons (**Fig. 6G**; **Extended Data Fig. 9J,L**). Together, these results demonstrate that interruption of the pattern generating network in HVC, by either optogenetic perturbations or silencing of HVC_X_ neurons, drives song truncation and likely releases a common circuit mechanism driving rapid restarting of the neural sequence for song.

## Discussion

The precise sequencing of neuronal activity in HVC has been proposed to function as a clock, which controls the timing and progression of zebra finch song syllables and inspiratory pauses between syllables^4,6–8,10^. Considerable debate has centered on whether these patterns of activity require instructive patterned input for song progression. Yet, pathway and cell-type selective manipulations required to directly address this debate have been lacking. Here, we provide key observations indicating that HVC functions as a sequence generating network that, in adult birds, does not require patterned input, at least from its best described afferent pathways, to achieve the patterned output necessary to complete the song motif.

Chunking of motor sequences followed by concatenation or fusing of commonly repeated sequences is a proposed mechanism for optimizing learning and performance^46–48^. Motor chunking is, for example, thought to function in learning and production of the movement sequences needed for fluent speech production, as well as other behaviors. Accordingly, early stages of juvenile bird’s song development have been shown to involve splitting and growth of neural sequences in HVC as new syllables are being learned^49^. This process likely reflects chunking of respiratory and vocal patterns needed for accurate and rapid learning. Juvenile songbirds progressively shape their song, practicing thousands of times per day, and pallial input pathways are necessary to direct such developmental song learning^50–52^. Here, we show that in adults with fully mature song these major pallial input pathways to HVC are dispensable for song production. Moreover, our cell-type selective manipulations in HVC demonstrate that the ordered syllable sequence of the song motif spontaneously restarts if it is prematurely truncated. This skipping of song back to the beginning is reminiscent of CPG rhythm resetting and consistent with the holistic control of the motif by a sequence-generating circuit in HVC^15,29^. These findings suggest that the phase at the end of song development, referred to as ‘crystallization’, involves consolidation of neural programs for motor control within HVC. We hypothesize that as song developmentally becomes more stereotyped and precise, consistent daily practice concatenates these sparse neural sequences into a stable chain that can autonomously sustain song motif completion.

Interhemispheric coordination is likely to be fundamental for proper song production^20,21^. Birds lack a corpus callosum, therefore our findings illustrating the autonomy of HVC from afferent inputs raises questions about how the interhemispheric timing of activity in sequence pattern generating circuits is coordinated. Uva receives bilateral ascending input from the respiratory medulla and midbrain vocal circuits and is therefore hypothesized to play a prominent role in interhemispheric coordination of HVC^9,10,18^. Accordingly, we find that thalamic input from Uva is permissive for initiating the song motif. However, our evidence indicates that it is not needed for transitioning from syllable-to-syllable within the motif. Uva may therefore send synchronized onset cues for song to coordinate initiation of each motif in the song bout, that then continues autonomously in each HVC. Nonetheless, Uva remains active throughout the motif^18^ and thus, could also function like a metronome that provides timing signals helping interhemispheric coordination without being required to instruct transitions within ongoing song motifs.

Our synaptic connectivity mapping identifies a prominent heterosynaptic connectivity pattern, with HVC_X_ neurons consistently making monosynaptic connections with HVC_RA_ neurons but only sparsely with other HVC_X_ neurons. Together with the previous literature, this supports the idea that the major synaptic connectivity within HVC involves disynaptic inhibition and monosynaptic excitation between HVC_RA_ and HVC_X_ neurons, rather than homotypic connections within either class of excitatory neurons^41,42^. More research will be needed to fully describe cell-type connectivity in the network and understand how song sequence progression is fully controlled. For example, HVC contains a sparse population of excitatory neurons that project to the auditory system^53^. Additionally, HVC receives a sparse input from RA^54,55^. The role of these pathways in song motor control is less well understood and not addressed in the current research. Nonetheless, we propose here a straightforward computational model of HVC inspired by our present results that can sustain sequence generation and song restarting following circuit perturbations. Moreover, model simulations indicate that substantial reduction of HVC_X_ neuronal transmission leads to stochastic song truncations and restarting of the motif – a prediction matched by our selective expression of TeNT in these neurons. However, the HVC song circuit appears to be robust to moderate perturbation, as shown by the lack of effect of previous HVC_X_ ablation studies^39,40^, focal lesions studies^56^, our model predictions (**Fig. 6E**), and seen in our own data where birds with lower levels of TeNT expression did not exhibit significant disruptions in song (**Fig. 6H,I**; **Extended Data Fig. 9G-I**).

In summary, this research reveals that a premotor circuit can holistically control strings of vocal syllables initiated by thalamic input and it has helped better define the synaptic circuit architecture critical to this pattern generating circuit. The key insights of this research emerged through cell-type and pathway selective manipulations of neural circuits in freely singing birds. In future experiments it will be important to examine how ‘fused’ sequence elements are integrated for the control of even more complex behaviors. Zebra finches produce only a single stereotyped sequence of song syllables, making them an ideal model for first testing how the brain controls strings of vocal syllables. Other songbird species learn larger repertoires of songs and/or string together short sequences of song syllables in more complex ways. We hypothesize that chunking, followed by concatenation of reliably reproduced neuronal sequences, underlies these behaviors and that the approaches applied here can help identify the boundaries of the motor sequences used by the brain to support production of complex vocal behaviors. It is likely that many other naturally well-learned sequenced behaviors (e.g. human speech articulation, rodent oromanual food manipulation) are also supported by similar circuit mechanisms. Demonstrating holistic cortical control of motor sequences in the songbird further suggests that this could be an evolutionarily conserved strategy for learning and controlling sequenced behaviors.

---

**Extended Data Fig.1.**
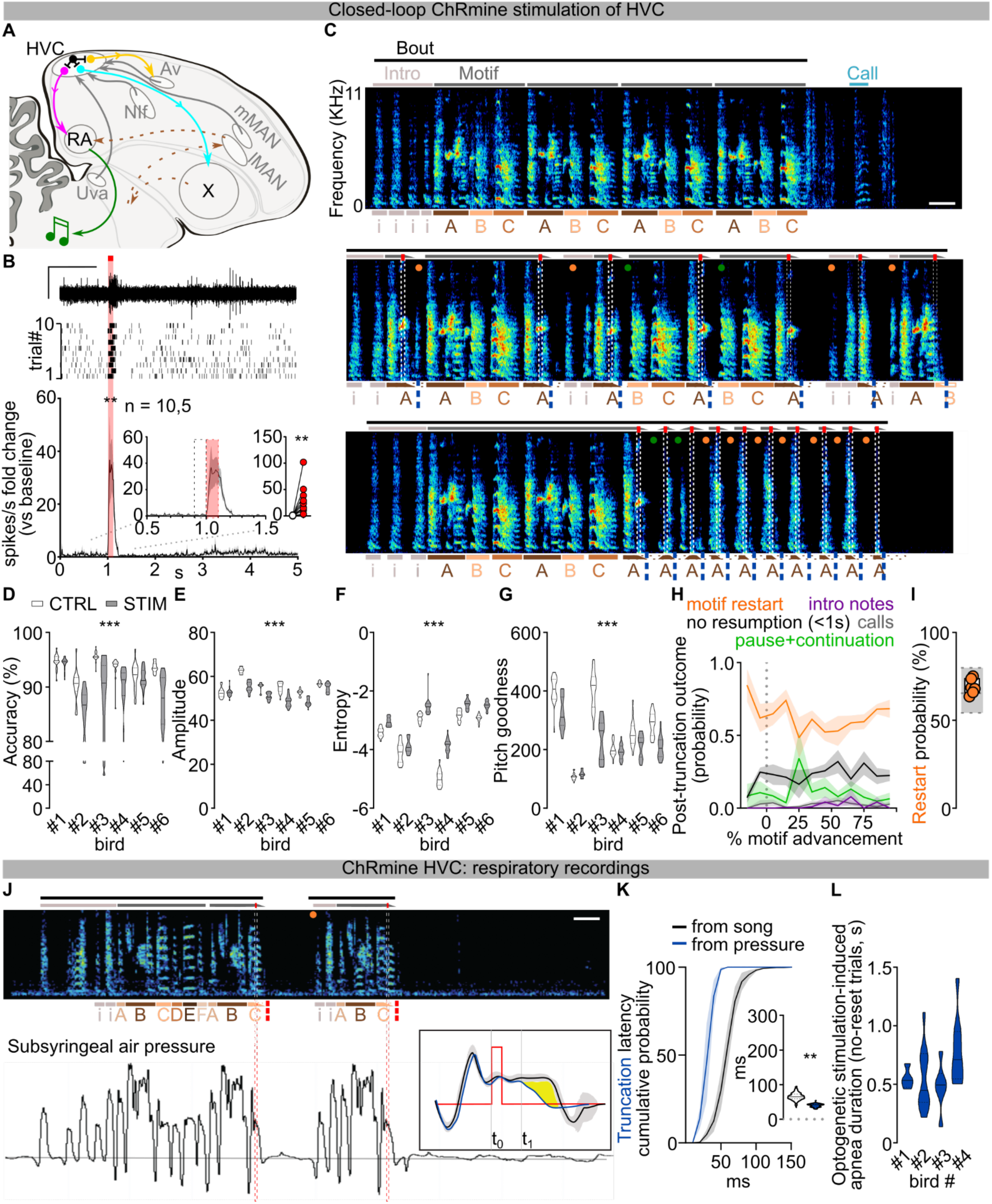
Effects of optogenetic stimulation of HVC neurons in singing zebra finches. Schematic of zebra finch song circuits (sagittal view), including HVC’s afferents (grey) from Uva (nucleus Uvaeformis), NIf (nucleus interface of the nidopallium), mMAN (medial magnocellular nucleus of the anterior nidopallium), and Av (nucleus avalanche); HVC projections to RA (robust nucleus of the arcopallium) via HVC_RA_ neurons (magenta), to the striatopallidal region Area X via HVC_X_ neurons (cyan), and to Av via HVCAv neurons (yellow); HVC inhibitory interneurons (black); the cortico-basal ganglia-thalamocortical song pathway (brown dashed lines), and the corticobulbar song motor pathway from RA (green). **B)** HVC multiunit neuronal activity recording in anesthetized birds expressing ChRmine in HVC. Sample trace (top, scalebar 1s, 1V), raster plot (mid, 10 trials) and normalized peri-stimulus time histogram (bottom) reporting the change in multi-unit HVC firing activity in response to light stimulation (100ms, red bar; two-way ANOVA, F(1,9)=11.20, P=0.0086); inset displays magnified detail of the PSTH and scatter plot highlighting and computing the average (per hemisphere) response to the first 100ms light stimulation (red dashed rectangle) compared to the last 100ms baseline (black dashed rectangle, Wilcoxon test, P=0.002; n= 10 hemispheres, 5 birds). **C)** More spectrograms from the bird in Fig. 1B displaying multiple events of optogenetically-evoked truncations followed by rapid restart of a motif (orange circle) or continuation of the motif after a pause (green) normally not present in the unstimulated motif within 1 second following stimulation. **D)** Violin plots reporting accuracy of song segments with (gray) and without (white) stimulation for each bird (n=6; two-way ANOVA, CTRL vs STIM, F(1,114)=55.22, P<0.001). **E-G)** Same as **D** but for Amplitude (two-way ANOVA, CTRL vs STIM, F(1,114)=246.4, P<0.001), Entropy (two-way ANOVA, CTRL vs STIM, F(1,114)=322.5, P<0.001) and Goodness of pitch (two-way ANOVA, CTRL vs STIM, F(1,114)=101.4, P<0.001). **H)** Average ±SEM probability of post-truncation behavior (no vocalization resumption in the 1s post-truncation (black), motif restart with any introductory note or syllable A (orange), intro notes not followed by a motif (purple), calls (grey), resumption of the motif after a pause (green)) following HVC light stimulation delivered at different latencies throughout the motif (% motif advancement, bins = 10% motif advancement). **I)** Box and scatter plot reporting the probability of motif restart for each bird (orange dots). The underlying gray box shows the distribution of each of the birds’ probability of producing a motif after any one motif (see methods), providing a basis for normalization of motif restart probability (gray box reports maximum, minimum, and median of the 6 birds). **J)** Spectrogram (top) and subsyringeal air pressure (bottom, black trace) relative to 3 motifs (marked by gray bars above the spectrogram, stimulation: red bars, 10ms, lettering and symbols as per panel B, scalebar 200ms). (bottom) Grey semitransparent horizontal line indicates ambient pressure, supratmospheric pressure shows expiratory air pressure and subatmospheric is inspiratory. The insert (right) shows the average of syllable C (grey shading ±2 SD) in control (black) and stimulated trials (blue line). The yellow shading indicates the significant reduction in air pressure caused by optogenetic stimulation HVC neurons. **K)** Average ±SEM of cumulative probability distributions calculated for each bird whose pressure was recorded, displaying the latency to truncation as measured for each bird from spectrograms (black line) or from subsyringeal pressure (blue line) (10ms time bins, two-way ANOVA, F(1,3)=19.50, P<0.001, Dunnett’s post-hoc identifies significant difference (p<0.05) between 40 and 60ms time bins); (inset) violin plots reporting the latency of motif truncation computed across all the birds (latency calculated from spectrograms (white), latency calculated from pressure for the same 4 birds (blue); Mann-Whitney U=236, P<0.001). **L)** Violin plots reporting the duration of the optogenetically-evoked apnea in events where truncations were not followed by restarting the song motif, computed across all stimulations for each of the 4 birds.

**Extended Data Fig.2.**
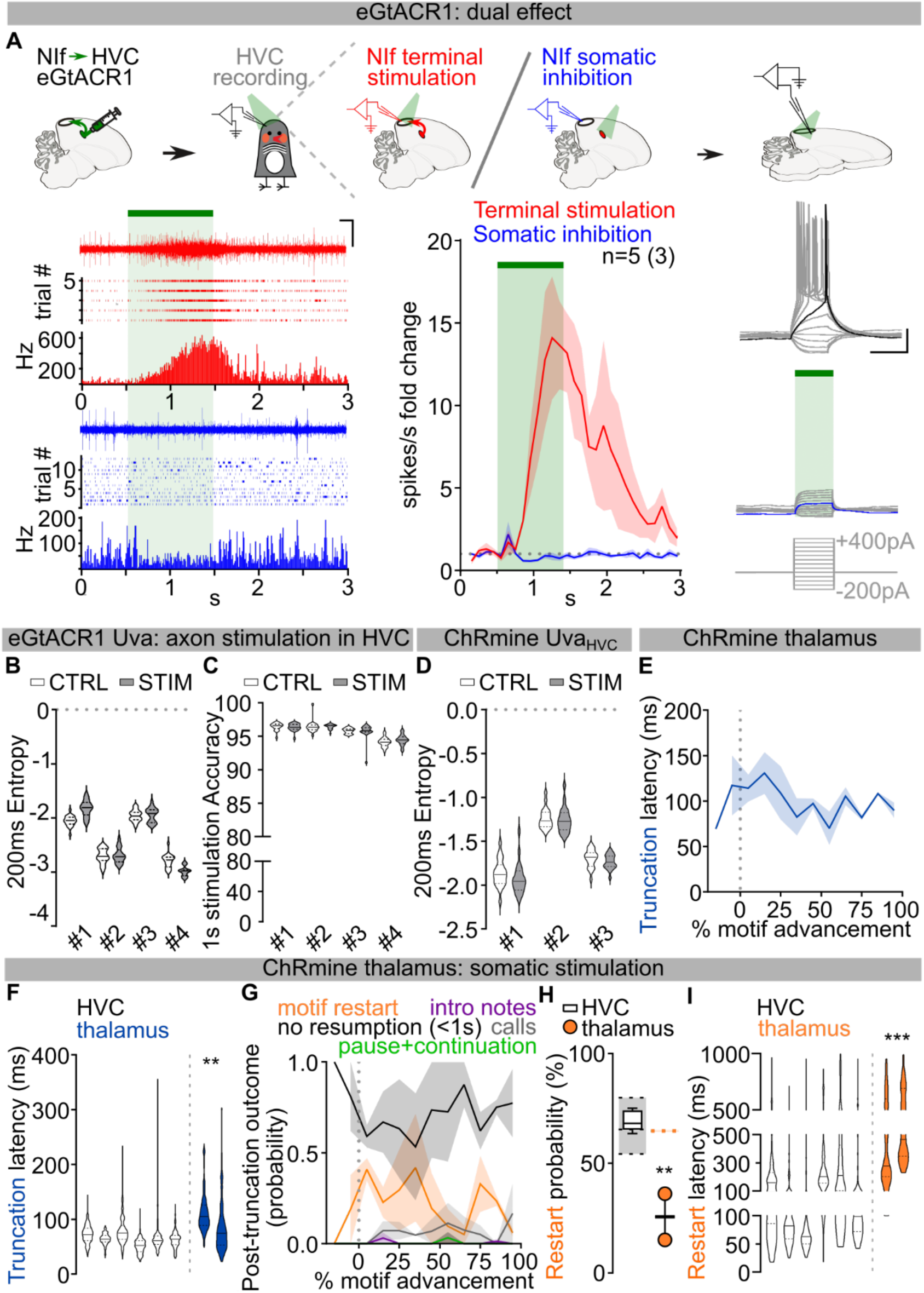
Role of Uva and Peri-Uva thalamus in song. **A)** Schematics (top), and raster plots and sample traces (bottom) showing the dual effect of eGtACR1-mediated stimulation of axonal terminals causing neurotransmitter release (schematized in red), and somatic inhibition (blue). (left) Sample traces, raster plots and PSTH reporting in-vivo recordings of HVC neuronal activity change in response to NIf axon terminal excitation (red) or soma inhibition (blue) upon light delivery (region shaded in green; scalebar 200ms, 1V). (center) Average ±SEM fold change in HVC activity upon NIf axonal stimulation (red) or somatic inhibition (blue) (binned each 100ms). (right) Sample traces of evoked EPSPs from an eGtACR1-expressing neuron in absence or presence of somatic light-mediated inhibition (green bar; current injection 100ms 50pA steps, −200pA to +400pA, scalebar 100ms, 20mV). **B,C)** Data from birds expressing eGtACR1 in Uva and implanted with optic fibers over HVC. B) Violin plots reporting Entropy of song segments with (gray) and without (white) stimulation (n=4); two-way ANOVA, CTRL vs STIM, F(1,76) = 1.099, P=0.2979). C) Violin plots reporting accuracy of song segments with (gray) and without (white) stimulation (n=4) when using 1 second long stimulation (two-way ANOVA, CTRL vs STIM, F(1,76)=0.06636, P=0.7974). **D)** Data from birds expressing ChRmine in UvaHVC neurons and implanted with optic fibers over Uva. Violin plots reporting Entropy of song segments with (200ms stimulation (gray)) and without (white) stimulation (n=4; two-way ANOVA, CTRL vs STIM, F(1,57)=3.500, P=0.0665). **E)** Average latency ±SEM of motif truncation in response to thalamic light stimulation across the motif (% motif advancement, bins = 10% motif advancement). **F)** Violin plots reporting the latency of motif truncation upon light stimulation, per bird (thalamus stimulation (blue), HVC stimulation (white, dataset from Fig.1); nested one-way ANOVA comparing all datasets across the manuscript, F(6,19)=9.678, P<0.001, Dunnett’s post-hoc P=0.0024). **G)** Average ±SEM probability of post-truncation behavior following thalamic light stimulation (no vocalization resumption in the 1s post-truncation (black), motif restart with any introductory note or syllable A (orange), intro notes not followed by a motif (purple), calls (grey), resumption of the motif after a pause (green)) across the motif (% motif advancement, bins = 10% motif advancement). **H)** Box and scatter plot reporting the probability of motif restart (thalamus-stimulated birds (orange dots); empty box plot representing data from birds receiving HVC stimulation reported from Extended Data Fig. 1I; one-way ANOVA, F(5,16)=5.717, P=0.0033, Dunnett’s post-hoc P=0.0014). The underlying gray box shows the probability of producing a motif after any one motif (see methods), providing a basis for normalization of motif restart probability (gray box reports maximum, minimum, and median). **I)** Violin plots reporting the latency of motif restart (thalamus stimulation (orange), white: HVC stimulation dataset from Fig.1 compared against all experimental groups across the manuscript; nested one-way ANOVA, (5,16)=5.674, P=0.0034, Dunn’s post-hoc P<0.001).

**Extended Data Fig.3.**
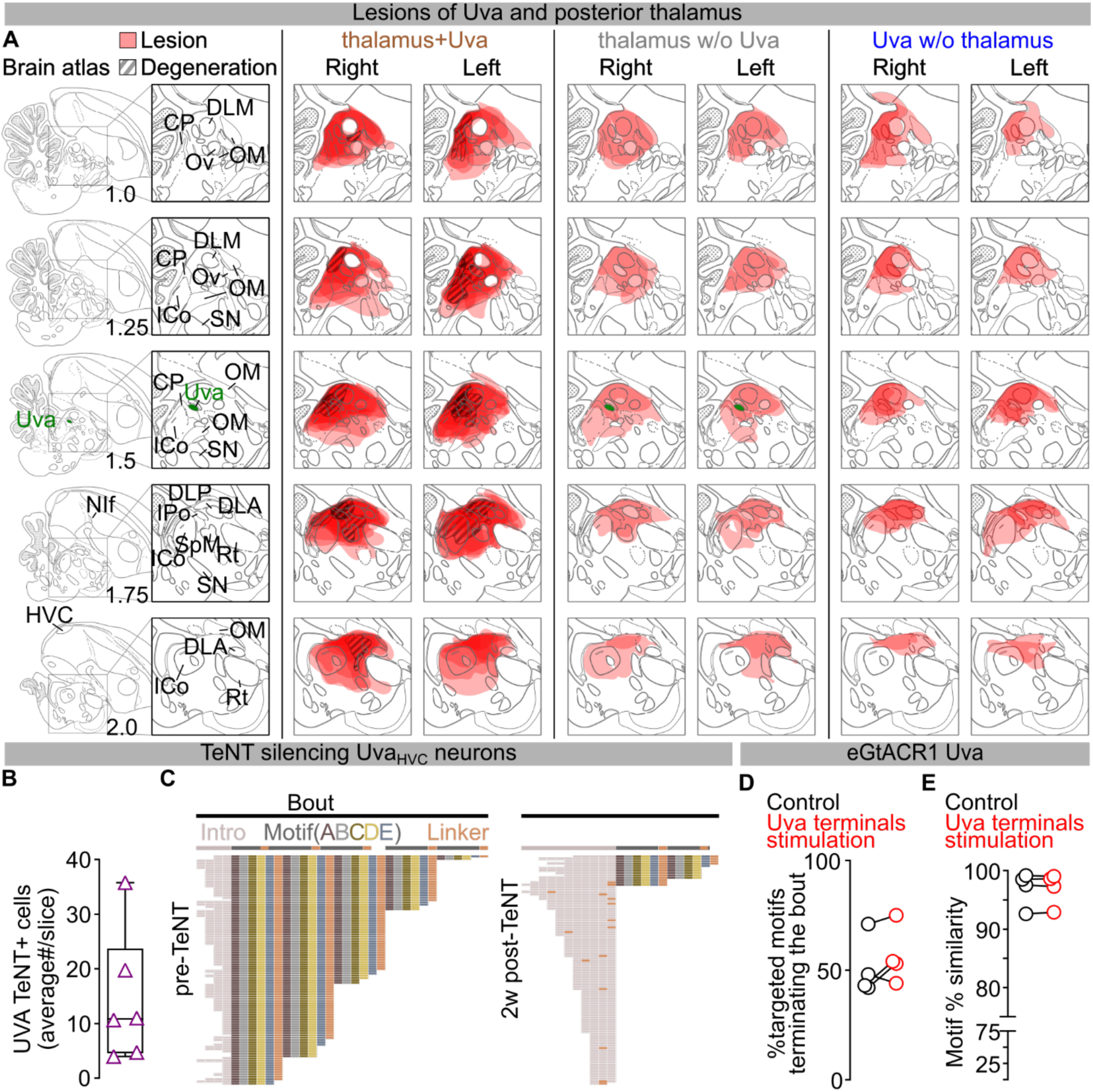
Role of Uva in motif initiation. **A)** Mapping of excitotoxic thalamic lesions. Brain atlas plates adapted from the zebra finch atlas (http://www.zebrafinchatlas.org), from 1 to 2mm across the mediolateral axis of magnified of the peri-Uva thalamic area (Uva highlighted in green). Extent of the lesion as measured from the lack of NeuN staining and divided by subgroup (peri- thalamus+Uva (left), peri-thalamus excluding Uva (middle), Uva excluding the larger perithalamic areas (right)). DLM – medial part of the dorsolateral nucleus of the anterior thalamus, CP – posterior commissure, Ov – nucleus ovoidalis, OM – occipitomesencephalic tract, ICo – intercollicular nucleus, SN – substantia nigra, Uva – nucleus Uvaeformis, DLP – dorsolateral nucleus of the posterior thalamus, DLA – dorsolateral nucleus of the anterior thalamus, IPo – intermedioposterior nucleus, SpM – nucelus spiriformis medialis, Rt – nucleus rotundus, NIf – nucleus interfacialis). **B)** Average number of TeNT expressing UvaHVC neurons/brain slice (purple, n=6 birds). **C)** Syntax raster plots (∼100 song bouts/day) from bird in Fig.3F showing syntax changes due to TeNT expression in UvaHVC neurons. **D,E)** Optical excitation of Uva terminals over HVC in the birds from Fig. 3K doesn’t significantly affect song bout terminations (D) (n=4 birds, control :black, no light, Uva terminals excitation: red, light delivered over eGtACR1-expressing Uva neurons for 1 sec; paired t-test, P=0.2211) nor motif self-similarity (E) (paired t-test, P=0.9429).

**Extended Data Fig.4.**
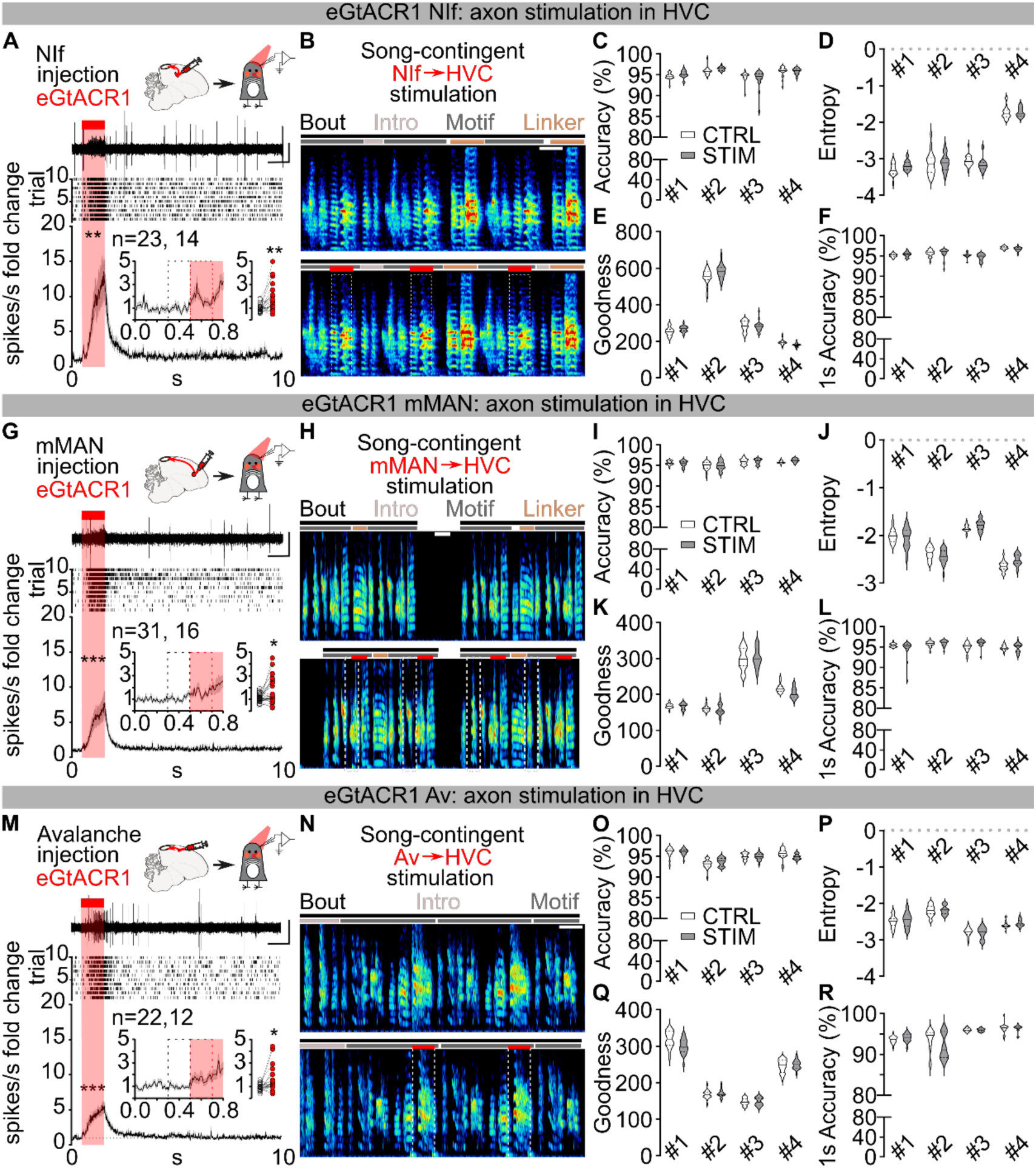
Optogenetic stimulation of pallial afferents to HVC does not disrupt the song motif. **A)** Schematic of in-vivo recording of HVC multiunit neuronal activity in anesthetized birds expressing eGtACR1 in NIf; sample trace (top), raster plot (mid, 10 trials) and normalized peri-stimulus time histogram (bottom) reporting the change in multi-unit HVC firing activity in response to light stimulation of eGtACR1-expressing NIf afferents (1s, red bar; two-way ANOVA, F(1,22)=12.07, P=0.0022). Inset shows magnified PSTH and scatter plot with the average (per hemisphere) response to the first 200ms light stimulation (red dashed rectangle) compared to the last 200ms baseline (black dashed rectangle, Wilcoxon test, P=0.0017; n= hemispheres, birds). **B)** (top) Schematic of song-contingent light stimulation of NIf axonal terminals in HVC (optic fiber implanted over HVC) and sample spectrogram of unstimulated (top) and stimulated song (bottom, red bars, 200ms ≈10mW bilateral red LED, spectrogram scale, 0-11KHz, scalebar 200ms). **C)** Violin plots reporting accuracy of the stimulated song segment (gray), or corresponding control unstimulated segment (white) per each bird (n=4); two-way ANOVA, CTRL vs STIM, F(1,76) = 0.7208, P=0.3985). **D)** Same as C but for Entropy (n=4; two-way ANOVA, CTRL vs STIM, F(1,76) = 0.1807, P=0.6720). **E)** same as C but for goodness of pitch (n=4; two-way ANOVA, CTRL vs STIM, F(1,76) = 2.301, P=0.1334). **F)** same as C but for Accuracy of the entire motif for birds receiving 1s light stimulation (n=4; two-way ANOVA, CTRL vs STIM, F(1,76) = 3.072, P=0.0837). **G-L)** same as A-F, but for eGtACR1 expression in mMAN (n=4; (n=4; I: two-way ANOVA, CTRL vs STIM, F(1,76) = 0.6008, P=0.4407; J: two-way ANOVA, CTRL vs STIM, F(1,76) = 0.1036, P=0.3119, K: two-way ANOVA, CTRL vs STIM, F(1,76) = 2.828, P=0.0967, L: two-way ANOVA, CTRL vs STIM, F(1,76) = 0.2599, P=0.6117). **M-R)** same as A-F, but for eGtACR1 expression in Av (n=4; O: two-way ANOVA, CTRL vs STIM, F(1,76) ) = 0.1304, P=0.7190O: two-way ANOVA, CTRL vs STIM, F(1,76) = 0.7229, P=0.3979, P: two-way ANOVA, CTRL vs STIM, F(1,76) = 0.1634, P=0.2050, Q: two-way ANOVA, CTRL vs STIM, F(1,76) = 1.143, P=0.2883).

**Extended Data Fig.5.**
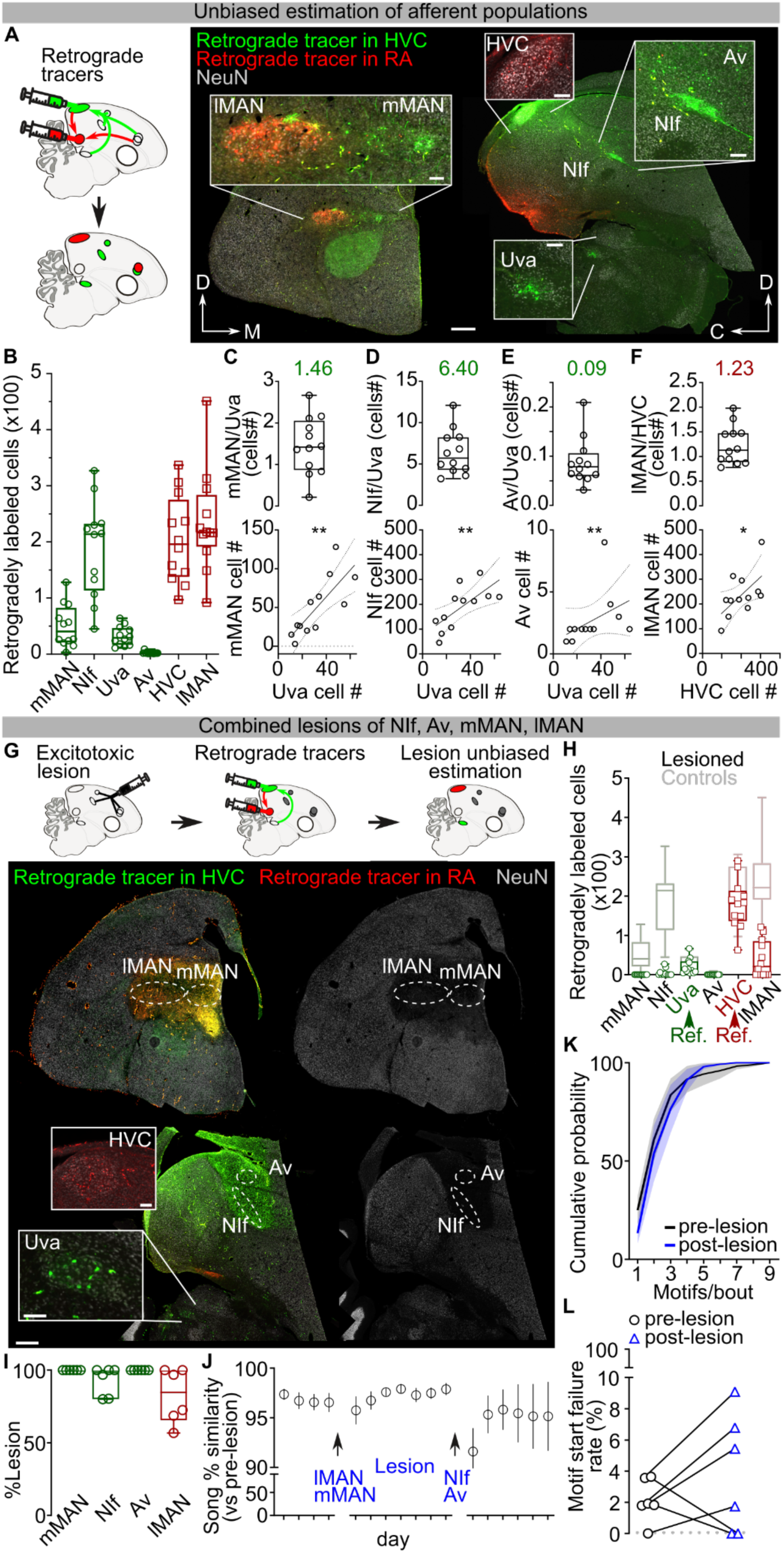
Concurrent lesions of pallial afferents to HVC and RA does not disrupt the song motif. **A)** Schematic and sample images of retrogradely labeled HVC afferent neurons in NIf, Uva, mMAN and Av (green) and RA afferent neurons in lMAN and HVC (red) (coronal slices for LMAN and mMAN, sagittal slices for HVC, NIf, Av, Uva; scalebar 500µm, insets 100µm). **B)** Box and scatter plot reporting the number of retrogradely labeled neurons in each brain area per hemisphere (green projecting to HVC, red to RA, n=12 hemispheres, 7 birds). **C)** box and scatter plot (top) and correlation (bottom) reporting the ratio of the number of retrogradely labeled neurons in mMAN and Uva (average=1.46, R^2^=0.6124, Spearman r=0.7972, P=0.0029). **D)** same as L but for the number of neurons in NIf and Uva (average=6.40, R^2^=0.4917, Spearman r=0.7902, P=0.0033). **E)** same as L but for the number of neurons in Av and Uva (average = 0.09, R^2^=0.1726, Spearman r=0.7112, P=0.0093). **F)** same as L but for the number of neurons in lMAN and HVC (average = 1.23, R^2^=0.3486, Spearman r=0.6713, P=0.0202). **G)** Schematic and sample image of the excitotoxic lesion of lMAN, mMAN, NIf and Av, combined with retrograde labeling from tracer injections in HVC (green) and RA (red). Tracer labeling reveals surviving afferent neurons and allows post-hoc unbiased estimation of the lesion extent (insets display magnified detail of retrogradely labeled HVC_RA_ and UvaHVC cells; scalebar 500µm, insets 100µm). **H)** Box and scatter plot reporting the number of surviving retrogradely labeled neurons in each brain area per hemisphere (green projecting to HVC, red to RA, retrograde labeling in Uva and HVC are reference areas for unbiased lesion quantification, n=12, 6 birds). Grayed out box plots outlines from panel B reports control data for ease of comparison of the lesion extent. **I)** Quantification of mMAN, NIf, Av and lMAN lesion, per bird (n=6 birds). **J)** Time course of song self-similarity (avg ±SEM, One-Way ANOVA, Mixed-effects analysis, F (1.129, 5.364)=1.599, P=0.2640). **K)** Cumulative probability curves reporting the number of motifs/bout sung by the birds before (black) and after (blue) the bilateral excitotoxic lesion of NIf, Av, mMAN and lMAN (n=6 birds, two-way ANOVA, F(1,45)=0.6098, P=0.4390). **L)** Scatter plot of motif start failures before (black circles) and after (blue triangles) the bilateral excitotoxic lesion of NIf, Av, mMAN and lMAN (Wilcoxon test, P=0.4375).

**Extended Data Fig.6.**
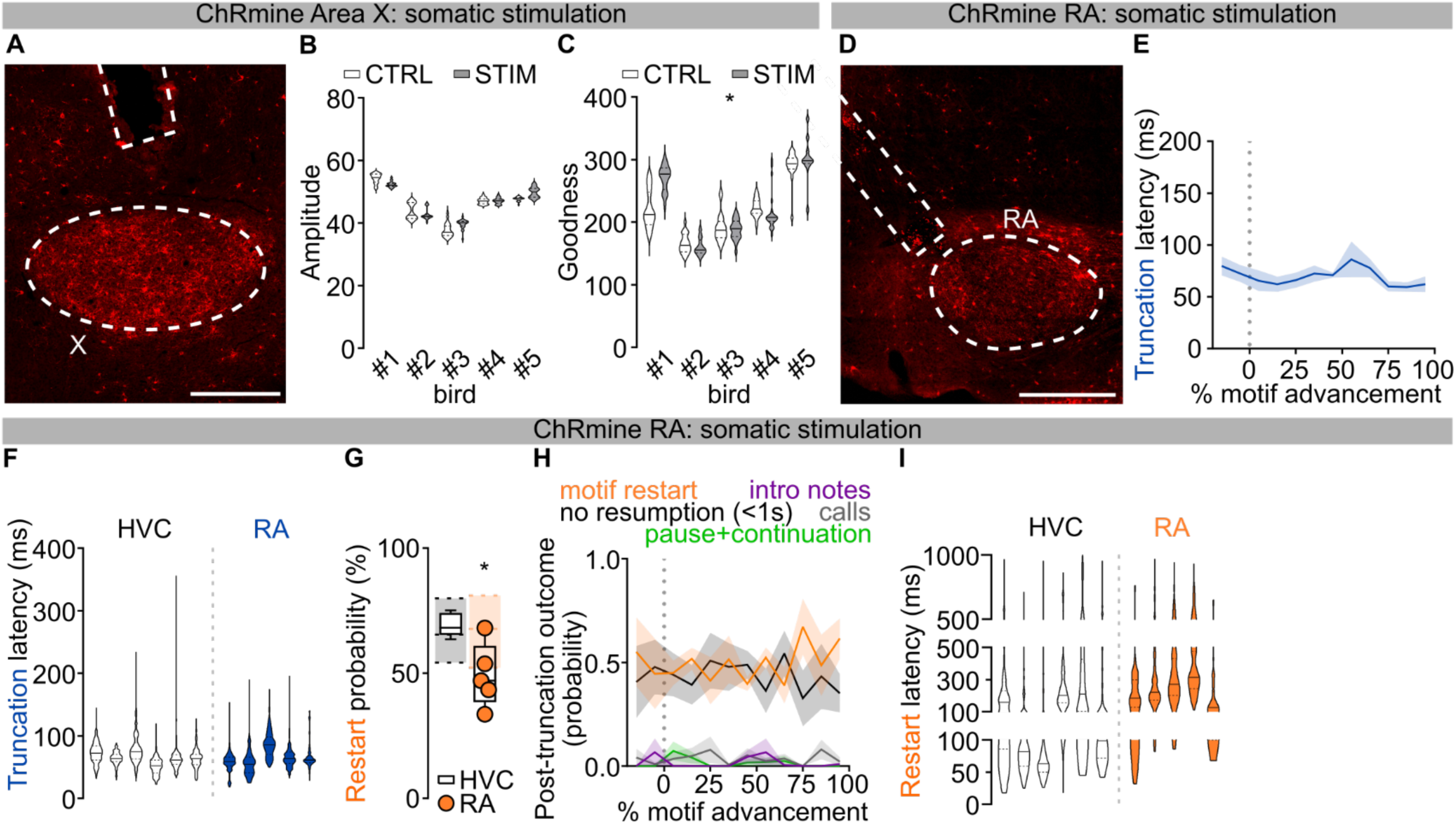
Optogenetic excitation of Area X and RA. **A)** ChRmine expression in Area X and outline of the optic fiber track (scalebar 500µm). **B-C)** Violin plots reporting amplitude and goodness of pitch of song segments with (gray) and without (white) Area X stimulation for each bird (n=5; amplitude: two-way ANOVA, CTRL vs STIM, F(1,114)=0.5747, P=0.4503; goodness of pitch: two-way ANOVA, F(1,114)=6.606, P=0.0117). **D)** ChRmine expression in RA and outline of the optic fiber track (scalebar 500µm). **E)** Average latency ±SEM of motif truncation in response to RA light stimulation (bins = 10% motif advancement). **F)** Latency to motif truncation following light stimulation of RA or HVC (blue: RA stimulation, white: HVC dataset from Fig.1; Nested one-way ANOVA comparing all datasets across the manuscript, F(6,19)=9.678, P<0.001, Dunnett’s post-hoc=0.9996). **G)** Probability of motif restart following optogenetic stimulation of RA or HVC (orange dots: RA-stimulated birds; empty box plot representing data from birds receiving HVC stimulation reported from Extended Data Fig. 1I. The underlying shaded areas represent the probability, for birds producing a motif after any one motif (see methods, provides the basis for normalization of motif restart probability; dashed lines maximum, median, and minimum); one-way ANOVA, F(5,16)=5.717, P=0.0033, Dunnett’s post-hoc P=0.04996). **H)** Average ±SEM probability of post-truncation behavior in response to the RA light stimulation delivered at different latencies throughout the progression of the motif(no vocalization resumption in the 1s post-truncation (black), motif restart with any introductory note or syllable A (orange), intro notes not followed by a motif (purple), calls (grey), resumption of the motif after a pause (green); bins = 10% motif advancement) I) Violin plots reporting the latency to motif restart (orange: RA stimulation birds, white: HVC stimulation dataset from Fig.1 compared against all experimental groups across the manuscript; Nested one-way ANOVA, (5,16)=5.674, P=0.0034, Dunn’s post-hoc P=0.2522).

**Extended Data Fig.7.**
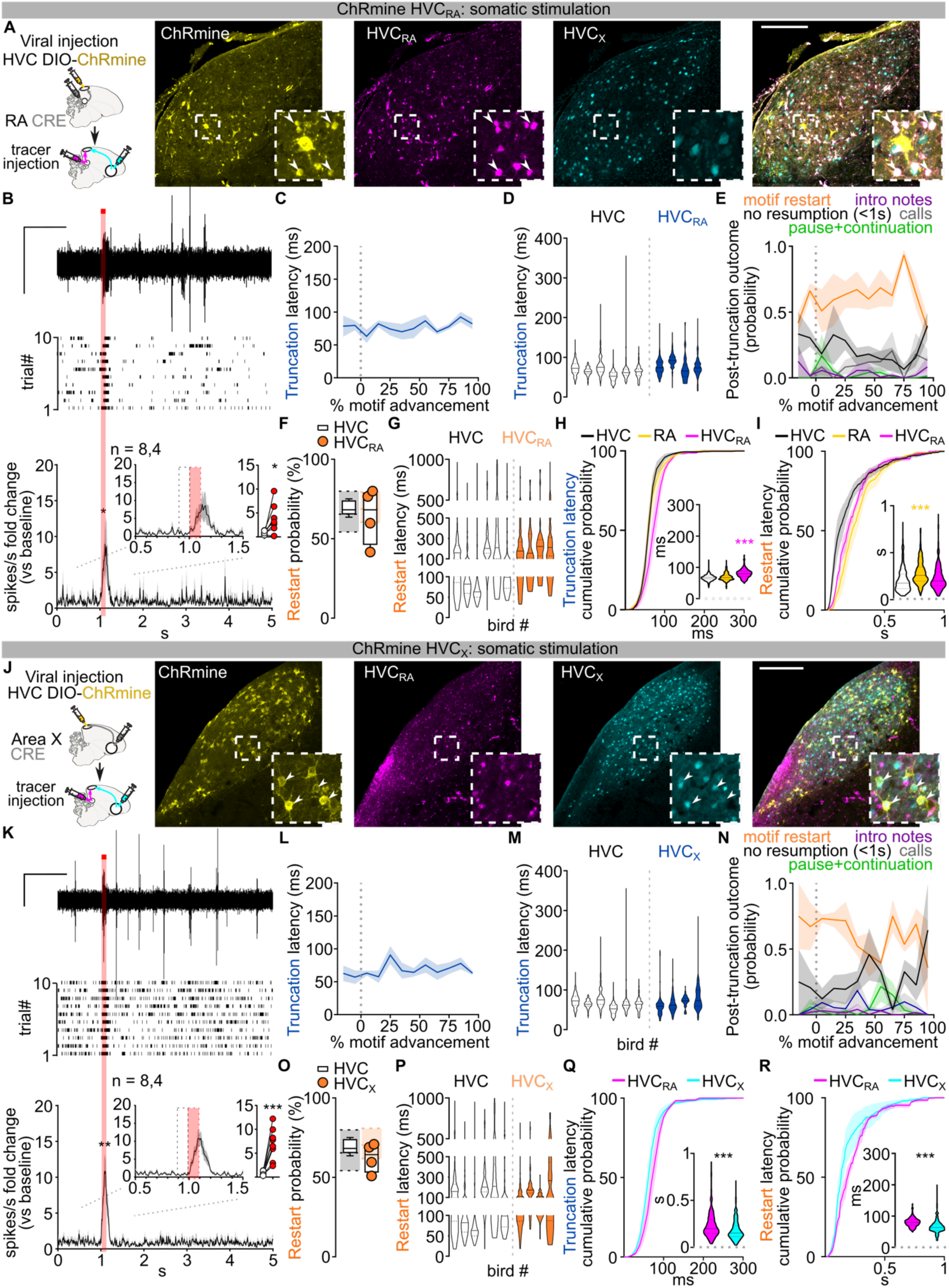
Optogenetic manipulation of HVC_RA_ and HVC_X_ neurons. **A)** Schematic of the viral strategy (AAV_DIO_ChRmine in HVC, high-titer retrograde AAV_Cre in RA) and sample images of retrogradely labeled HVC_RA_ (magenta, arrowheads) but not HVC_X_ neurons (cyan) displaying conditional expression of ChRmine (yellow) (scalebar 200µm, insets 3x magnification). **B)** HVC multiunit neuronal activity recording in anesthetized birds expressing ChRmine in HVC_RA_ neurons. Sample trace (top, scalebar 1V, 1s), raster plot (mid, 10 trials) and normalized peri-stimulus time histogram (bottom) reporting the change in multi-unit HVC firing activity in response to light stimulation (100ms, red bar; two-way ANOVA, F(1,7)=8.254, P=0.0239). (inset) PSTH and scatter plot illustrating the average (per hemisphere) response to the first 100ms light stimulation (red dashed rectangle) compared to the last 100ms baseline (black dashed rectangle, paired t-test, P=0.0178; n=8 hemispheres, 4 birds). **C)** Average latency ±SEM to motif truncation in response to the HVC_RA_ light stimulation (bins = 10% motif advancement). **D)** Latency to motif truncation following light stimulation (blue: HVC_RA_ stimulation, white: HVC dataset from Fig.1; nested one-way ANOVA comparing all datasets across the manuscript, F(6,19)=9.678, P<0.001, Dunnett’s post-hoc=0.3386). **E)** Average ±SEM probability of post-truncation behavior in response to the RA light stimulation computed based on the time of stimulation through the progression of the motif (no vocalization resumption in the 1s post-truncation (black), motif restart with any introductory note or syllable A (orange), intro notes not followed by a motif (purple), calls (grey), resumption of the motif after a pause (green); bins = 10% motif advancement). **F)** Probability of motif restart (orange dots: HVC_RA_-stimulated birds; empty box plot representing data from birds receiving HVC stimulation reported from Extended Data Fig. 1I. The underlying shaded areas represent the probability of producing a motif after any one motif (see methods, provides the basis for normalization of motif restart probability; dashed lines show the maximum, median, and minimum); one-way ANOVA, F(5,16)=5.717, P=0.0033, Dunnett’s post-hoc P=0.9963). **G)** Latency of motif restart (orange: HVC_RA_ stimulation birds, white: HVC stimulation dataset from Fig.1 compared against all experimental groups across the manuscript; nested one-way ANOVA, (5,16)=5.674, P=0.0034, Dunn’s post-hoc P=0.9236). **H-I)** Cumulative probability curves and violin plots (data reported from Fig. 4 and 5) illustrating the latency to song truncation (H) and latency to motif restart (I) in response to the light stimulation (average±SEM of each bird’s curve, magenta: HVC_RA_-stimulated birds, yellow: RA-stimulated birds, black: HVC-stimulated birds dataset from Fig.1 compared against all experimental groups across the manuscript; 10ms time bins). For truncation (H), two-way ANOVA, F(5,17)=4.134, P=0.0122, Dunnett’s post-hoc identifies significant difference (p<0.05) at the 60-90ms time bins; violin plots: one-way ANOVA, Kruskal Wallis test H(5)=467.6, post-hoc HVC vs. HVC_RA_ P<0.001). For restart latency (I) two-way ANOVA, F(6,22)=5.850, P<0.001, Dunnett’s post-hoc identifies significant difference (p<0.05) at the 170-200ms time bins; violin plots: one-way ANOVA, Kruskal Wallis test H(6)=245.5, post-hoc HVC vs. HVC_RA_ P=0.1157, HVC_RA_ vs. RA P<0.001). **J-P)** Same as A-G but for HVC_X_ neurons. K) 100ms light stimulation, red bar; two-way ANOVA, F(1,7)=23.52, P=0.0019); inset, paired t-test, P<0.001; n=8 hemispheres, 4 birds). M) Nested one-way ANOVA comparing all datasets across the manuscript, F(6,19)=9.678, P<0.001, Dunnett’s post-hoc=0.9782). O) One-way ANOVA, F(5,16)=5.717, P=0.0033, Dunnett’s post-hoc P=0.8772). P) Nested one-way ANOVA, (5,16)=5.674, P=0.0034, Dunn’s post-hoc P>0.9999). **Q)** Cumulative probability curves and violin plots (data reported from Fig. 5) illustrating the latency to song truncation in response to the light stimulation (magenta: HVC_RA_-stimulated birds, cyan: HVC_X_-stimulated birds; data compared across all the groups throughout the manuscript, 10ms time bins, two-way ANOVA, F(5,17)=4.134, P=0.0122, Dunnett’s post-hoc identifies significant difference (p<0.05) at the 60-80ms time bins; violin plots: one-way ANOVA, Kruskal Wallis test H(5)=467.6, post-hoc HVC_RA_ vs. HVC_X_ P<0.001). **R)** Same as Q but for latency to song restart (two-way ANOVA, F(6,22)=5.850, P<0.001, Dunnett’s post-hoc identifies significant difference (p<0.05) at the 170-200ms time bins; violin plots: One-Way ANOVA, Kruskal Wallis test H(6)=245.5, post-hoc HVC_RA_ vs. HVC_X_ P<0.001).

**Extended Data Fig.8.**
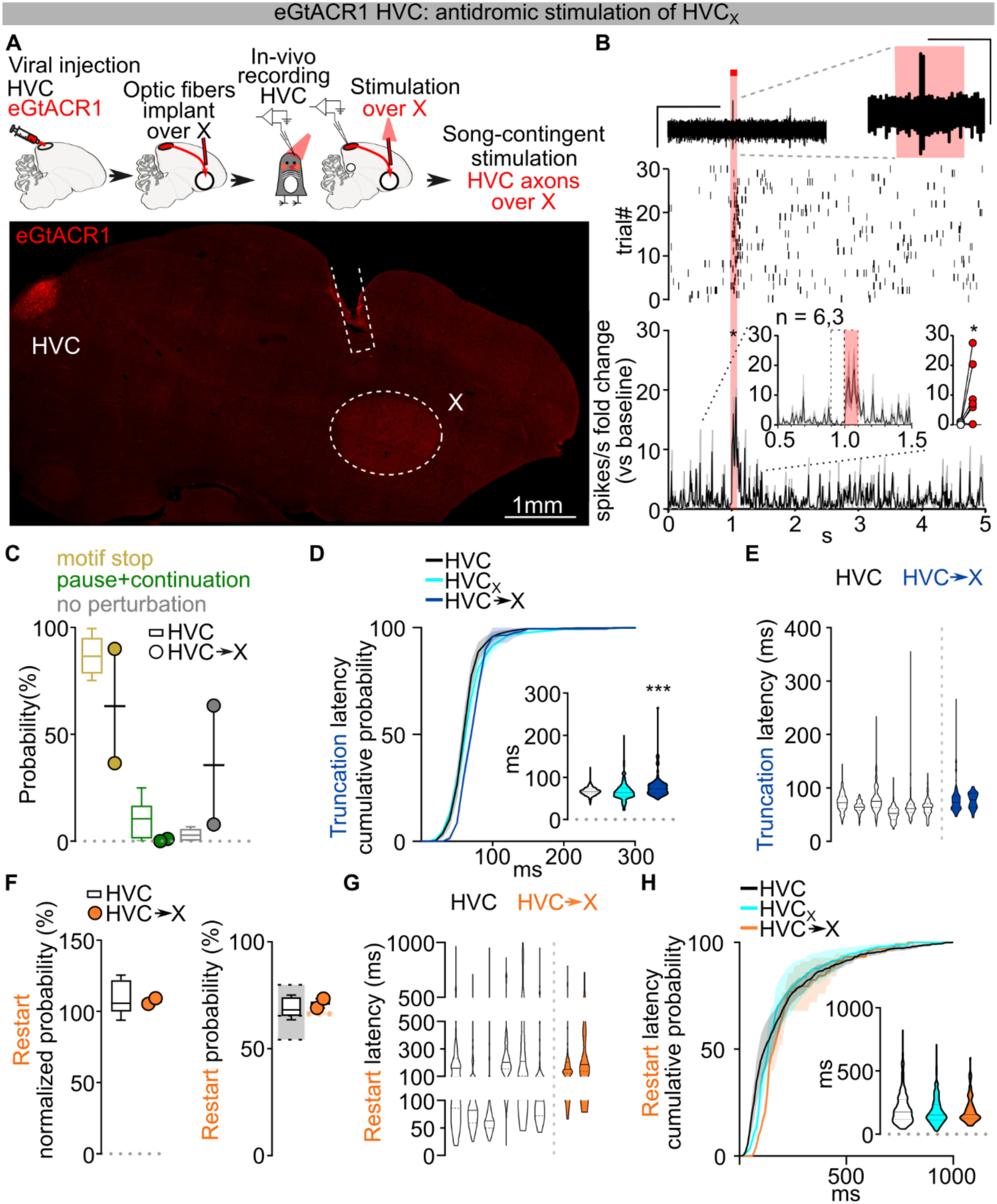
Antidromic optogenetic excitation of HVC_X_ neurons from Area X. **A)** Schematic and sample image of eGtACR1 expression in HVC and fiber optic implant over Area X (scalebar 1mm). **B)** HVC multiunit neuronal activity recording in anesthetized birds, sample trace (top, scalebar 1V, 1s; inset 1V, 100ms), raster plot (mid, 30 trials) and normalized peri-stimulus time histogram (bottom) reporting the change in multi-unit HVC firing activity in response to light stimulation of eGtACR1-expressing HVC_X_ afferents reaching Area X (100ms, blue bar). (inset) PSTH and scatter plot showing the average (per hemisphere) response to the first 100ms of light stimulation (red dashed rectangle) compared to the last 100ms baseline (black dashed rectangle, paired t-test, P=0.046; n= 6 hemispheres, 3 birds). **C)** Probability of motif truncation (okra), pause and continuation of the motif (green) or absence of syntactic perturbation (gray) after the light stimulation (HVC→X stimulated birds, n=2, filled circles; empty box plots from HVC stimulation in fig.1D reported for comparison; two-way ANOVA F(7,22)=2.549, P=0.0440, Dunnett’s post-hoc, motif stop P=0.0129, pause+continuation P=0.6958, no perturbation P<0.001). **D)** Cumulative probability curves reporting the latency to song truncation in response to the light stimulation (average ±SEM of each bird’s curve, blue: HVC→X, black: HVC, dataset from Fig.1 compared against all experimental groups across the manuscript, cyan: HVC_X_ dataset from Extended Data Fig. 7D for comparison,10ms time bins, two-way ANOVA, F(5,17)=4.134, P=0.0122, Dunnett’s post-hoc identifies significant difference (p<0.05) at the 50-80ms timebins). (inset) Latency of motif truncation computed across all the birds (blue: HVC→X, cyan: HVC_X_ dataset from Fig.6, white: HVC dataset from Fig.1 compared against all experimental groups across the manuscript; one-way ANOVA, Kruskal Wallis test, HVC vs. HVC→X P<0.001). **E)** Latency of motif truncation upon light stimulation, per bird (blue: HVC→X stimulation, white: HVC dataset from Fig.1; nested one-way ANOVA comparing all datasets across the manuscript, F(6,19)=9.678, P<0.001, Dunnett’s post-hoc=0.7105). **F)** Normalized (left) and not normalized (right) probability of motif restart for each bird (HVC→X stimulated birds: filled circles; empty box plots from Fig.1I and Extended Data Fig. 1I reported for comparison; for the not-normalized probability, the underlying shaded areas represent the probability, for each of the birds, of producing a motif after any one motif (see methods, provides the basis for normalization of motif restart probability; dashed top and bottom lines max and min, mid line represents median); left, one-way ANOVA, F(5,16)=9.217, P=0.0003, Dunnett’s post-hoc HVC vs. HVC→X: P>0.9999; right, one-way ANOVA, F(5,16)=5.717, P=0.0033, Dunnett’s post-hoc P=0.9996). **G)** Latency of motif restart (orange: HVC→X stimulation birds, white: HVC stimulation dataset from Fig.1 compared against all experimental groups across the manuscript; nested one-way ANOVA, (5,16)=5.674, P=0.0034, Dunn’s post-hoc P=0.9942). **H)** Cumulative probability curves reporting the latency to post-truncation motif restart in response to stimulation of HVC_X_ axon terminals (average ±SEM of each bird’s curve, orange: HVC→X, cyan: HVC_X_ dataset from Fig. 5, black: HVC dataset from Fig.1 compared against all experimental groups across the manuscript, 10ms time bins, 2W ANOVA, F(6,22)=5.850, P=0.0009, Dunnett’s post-hoc HVC→X vs. HVC identifies significant difference at the 80-90ms time bins; HVC→X vs. HVC_X_ P>0.05). (inset) Latency to motif restart computed across all the birds (orange: HVC→X, cyan: HVC_X_ dataset from Fig. 5, white: HVC birds dataset from Fig.1 compared against all experimental groups across the manuscript; one-way ANOVA, Kruskal Wallis test H(6)=245.5, post- hoc HVC→X vs. HVC_X_ P>0.9999).

**Extended Data Fig. 9.**
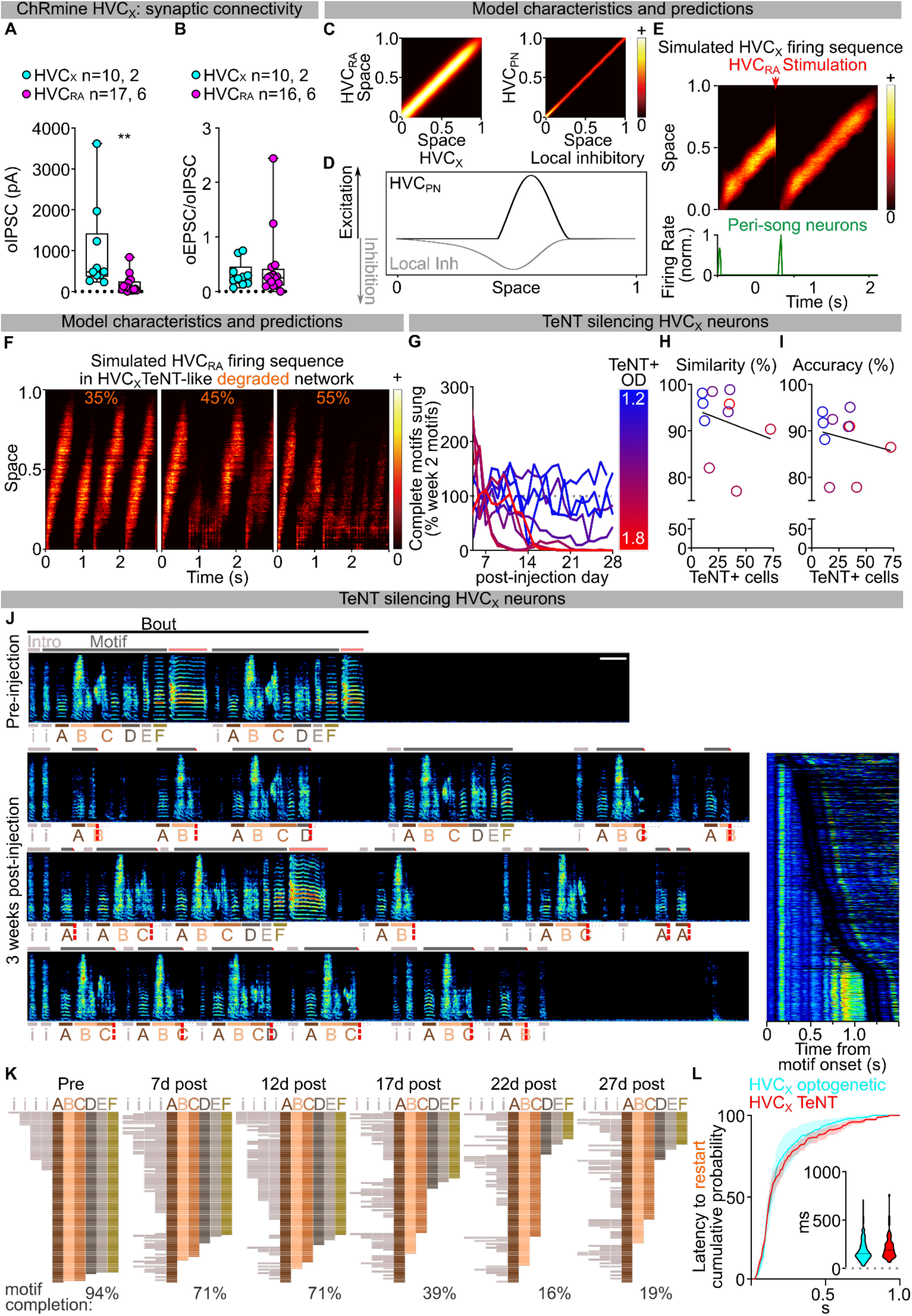
HVC_X_ neurons in song pattern generation. **A)** Box and scatter plot reporting oIPSC amplitude by cell class (Mann-Whitney test, P<0.001; n= cells, animals). **B)** Box and scatter plot of the ratio of oEPSC and oIPSC peak amplitude, per cell class (Mann-Whitney test, P=0.8159; n= cells, animals). **C)** Compensated connection matrix, (left) connection matrix between excitatory HVC_PN_ neuron classess; (right) connection matrix between excitatory and from local inhibitory neurons. **) D)** Excitatory (black) and inhibitory (gray) bumps are slightly offset so that that excitation in the network propagates in one direction. Firing rate is positive in both directions, downward curve for inhibitory neurons represents net inhibitory drive. **E)** Modeled/simulated HVC_X_ sequence displaying truncation and restart of the neuronal firing sequence following simulated optogenetic excitation of HVC_RA_ (red arrow). **F)** Simulated HVC_RA_ sequence displaying stochastic truncation in a degraded network mimicking the effect of TeNT expression in HVC_X_ neurons (reduced mean of the connection strength by 20% and took the variance sigma=0.5 (methods Eq. 5)). The neural activity will propagate along the chain utility broken by noise at random positions. **G)** Time course reporting data from Fig. 6H plotted by day, per bird. **H)** Scatter plot correlating the motif self-similarity (compared to the baseline motif) with the optical density of TeNT expression in HVC, per each bird (Spearman r=-0.2970, r^2^=0.05723, P=0.4069). **I)** Same as (H) but for the accuracy of the motif (Spearman r=-0.2848, r^2^=0.03963, P=0.4271). **J)** Spectrograms from a bird showing intermediate levels of TeNT expression in HVC_X_ neurons (different from the bird in Fig. 6). Notice the continuous failure to complete motifs, either with truncation within syllables or at syllable boundaries, followed by rapid motif restart (scalebar 200ms). The inset reports all spectrograms from one day in week 3, aligned at syllable A and ordered by motif length. **K)** Syntax raster plots (100 motifs/day) for the bird in panel (J). **L)** Cumulative probability curves reporting the latency to post-truncation motif restart (average ±SEM of each bird’s curve, blue: HVC_X_ TeNT, cyan: HVC_X_ dataset from Fig. 6, compared against all experimental groups across the manuscript, 10ms time bins, two-way ANOVA, F(6,22)=5.850, P=0.0009, Dunnett’s post-hoc HVC_X_ vs. HVC_X_ TeNT P>0.05). (inset) Violin plots reporting the latency of motif truncation computed across all the birds (HVC_X_ dataset from Fig. 5, HVC_X_ TeNT dataset from Fig. 6; one-way ANOVA, Kruskal Wallis test H(6)=245.5, post-hoc HVC_X_ vs. HVC_X_ TeNT P=0.5654).

## Supplementary video 1

3D rendering of zebra finch brain with tracers injected in HVC (green) and RA (red) to label efferent axons and retrogradely identified afferent neurons. RA axons flow caudally in the posterior commissure around and below Uva.

## Supplementary video 2

Animation displaying the model’s prediction of HVC_PN_ and interneuron activity waves in normal conditions, upon optogenetic stimulation, and in network degradation conditions mimicking TeNT expression in a subset of HVC_X_ neurons.

## Supplementary video 3

Animation of the schematic in Fig. 6B representing a proposed simplified description of HVC dynamics in normal conditions, upon optogenetic stimulation, and in network degradation conditions mimicking TeNT expression in a subset of HVC_X_ neurons.

## Materials and Methods

### Animals

Experiments described in this study were conducted using adult male zebra finches (120-500 days post hatch (dph)). All procedures were performed in accordance with protocols approved by Animal Care and Use Committee at UT Southwestern Medical Center.

### Viral Vectors

The following adeno-associated viral vectors were used in the experiments: rAAV2/9/fDIO-CBhCBh-eGTACR1- mScarlet, rAAV2/9/CBhCBh-Flippase, rAAV2/9/CBhCBh-ChRmine-mScarlet, rAAV2/9/DIO-CAG-ChRmine-mScarlet, rAAV2/9/DIO-CAG-TeNT-mScarlet (IDDRC Neuroconnectivity Core at Baylor College of Medicine), rAAV2/9/CMV-CRE-eGFP (Addgene). All viral vectors were aliquoted and stored at −80°C until use.

### Stereotaxic Surgery

Aseptic stereotaxic surgeries were performed after birds were anaesthetized (isoflurane inhalation, 0.8-1.5%). Viral injections were performed using previously described procedures^52,53^. Briefly, a cocktail of adeno-associated viral vectors (rAAV/CBhCBh-FLP in HVC, RA, Area X or Uva (2µl per hemisphere); 1:2 of rAAV/CBhCBh-FLP and rAAV/DIO-CBhCBh-eGtACR1 respectively (1-2µl total per hemisphere); rAAV/DIO-CAG-ChRmine in HVC or Uva (2µl) and rAAV/CMV-Cre in RA, Area X or HVC (0.5-1 and 2ul respectively); rAAV/DIO-Casp3 in Uva (2ul) and rAAV/CMV-CRE in HVC (2µl) were injected (1nl/s) into target areas with a Nanoject III (Drummond) and glass capillaries. Experiments were conducted starting a minimum of 3 weeks after viral injections. Fluorophore-conjugated retrograde tracers (Dextran 10,000MW, AlexaFluor 488, 568 and 647, Invitrogen; FastBlue, Polisciences) were injected bilaterally into Area X, Av, RA or HVC (160nl, 5x32 n, 32nl/s every 30 s) ^52,53,57^. Electrophysiologic mapping was used to determine the centers of HVC, NIf, mMAN, lMAN and RA, and Area X, Av and Uva were identified using stereotaxic coordinates (coordinates relative to interaural zero: head angle, rostral-caudal, medial-lateral, dorsal-ventral (in mm). HVC: 45°, AP 0, ML ±2.4, DV -0.2-0.6; NIf: 45°, AP 1.75, ML ± 1.75, DV −2.4 −1.8; mMAN: 20°, AP 5.1, ML ±0.6, DV −2.1 −1.6; lMAN: 20°, AP 5.1, ML ±1.7, DV −2.2 −1.6; RA: 70°, AP −1.5, ML ± 2.5, DV −2.4 −1.8; X: 45°, AP 4.6, ML ±1.6, DV -3.3 −2.7; Av: 45°, AP 1.65, ML ± 2.0, DV - 0.9; UVA: 20°, AP 2.5, ML ±1.6, DV -4.8 -4.2).

### Optogenetic Manipulations

For optogenetic stimulation, optic fibers (multimode 400µm, 0.39NA ThorLabs), were implanted bilaterally over HVC, RA, Area X or Uva using acrylic glue and dental cement. After recovery, the implanted fibers were connected to optic fibers through ceramic sleeves. The fibers were connected to a rotary joint and interfaced with a 1.5mm multimode fiber connected to an LED box (Prizmatix). Light intensity was regulated to achieve a final output of ≈10mW. To deliver optogenetic stimulation during song (200ms or 1s for HVC afferent stimulation, 10-50ms for direct ChRmine somatic stimulation or antidromic HVC_X_ stimulation), we used a custom software (pcaf^58^, LabVIEW). Random delays were introduced via the TTL output in 25ms progressive intervals. The entirety of the motif was targeted with successive trigger delay times.

#### Lesion Quantification

Excitotoxic lesion was induced by 1% ibotenic acid (50-100nl/injection site) or a cocktail of 1% ibotenic acid and 100mM Quisqualic acid (Uva and LMAN). Lesion extent was first verified by the absence or sparseness of NeuN immunostaining in the targeted nuclei. To provide an unbiased estimate of the lesion extent, retrograde tracers were injected in HVC and RA to highlight any surviving cells in the afferent nuclei. In control animals, the number of retrograde tracer-filled cells in each nucleus was quantified and correlations were calculated between cell counts in each nucleus (Supplementary Fig. 5A-F). This analysis provided a statistical validation to extrapolate the number of cells in a target nucleus from the number of cells counted in a reference nucleus. Therefore, an average ratio between each nucleus’ cell count was calculated. Based on these control ratios and on the number of cells in a non-lesioned reference nucleus, the expected number of retrogradely filled cells in each nucleus of each hemisphere was estimated.

### *In-vivo* extracellular recordings

To test the functional expression of opsins, we performed extracellular recording of HVC activity in birds under light isoflurane anesthesia (0.8%) with Carbostar carbon electrodes (impedance: 1670 µΩ-cm; Kation Scientific). A 400µm multimodal optical fiber was placed onthe brain surface overlaying HVC and delivered light stimulation (470nm, ≈20mW, 1s) during neural recordings. To test antidromic excitation of HVC_X_ neurons by axon terminal optical stimulation, optic fibers were implanted over Area X (470nm, ≈20mW, 100ms). Signals were acquired at 10kHz, and filtered (high-pass 300Hz, low-pass 20kHz). Spike rate (binned every 10ms) and peri-stimulus time histograms were calculated to quantify light stimulation responses (1-5 sites/hemisphere, Spike 2, Cambridge, EN). Birds without optically evoked responses were excluded from experiments. Spike counts and peri-stimulus time histograms were normalized to the pre-stimulus baseline (500ms). Two-way ANOVAs were calculated comparing the time course between stimulated and not stimulated recordings: for testing HVC afferents (1s stimulation), 0-5s (light stimulation 0.5-1.5s) vs. 5-10s (control, no stimulation); for ChRmine-expressing HVC neurons or HVC→Area X stimulation (100ms stimulation) 0.7-1.4s (300ms before and after 100ms light stimulation) vs. 5.7-6.4s (control, no stimulation). Wilcoxon tests ran on the average of the time course (intervals in the figure legends).

### *Ex vivo* physiology

#### Slice preparation

Zebra finches were deeply anesthetized, decapitated. The brain was removed from the skull and submerged in cold (1– 4°C) oxygenated dissection buffer. Acute sagittal 230 μm brain slices were cut in ice-cold carbogenated (95% O2/5% CO2) solution, containing (in mM) 110 choline chloride, 25 glucose, 25 NaHCO3, 7 MgCl2, 11.6 ascorbic acid, 3.1 sodium pyruvate, 2.5 KCl, 1.25 NaH2PO4, 0.5 CaCl2; 320-330 mOsm. Individual slices were incubated in a custom-made holding chamber filled with artificial cerebrospinal fluid (aCSF), containing (in mM): 126 NaCl, 3 KCl, 1.25 NaH2PO4, 26 NaHCO3, 10 D-(+)-glucose, 2 MgSO4, 2 CaCl2, 310mOsm, pH 7.3–7.4, aerated with a 95% O2/5% CO2 mix. Slices were incubated at 36 °C for 20 minutes, and then kept at room temperature for a minimum of 45 minutes before recordings.

#### Slice electrophysiological recording

Slices were constantly perfused in a submersion chamber with 32°C oxygenated normal aCSF. Patch pipettes were pulled to a final resistance of 3-5 MΩ from filamented borosilicate glass on a Sutter P-1000 horizontal puller. HVCPN classes, as identified by retrograde tracers, were visualized by epifluorescence imaging using a water immersion objective (×40, 0.8 numerical aperture) on an upright Olympus BX51 WI microscope, with video-assisted infrared CCD camera (Q-Imaging Rolera). Data were low-pass filtered (10 kHz) and acquired (10 kHz) (Axon MultiClamp 700B amplifier, Axon Digidata 1550B data acquisition, Clampex 10.6 Molecular Devices).

For voltage clamp whole-cell recordings, the internal solution contained (in mM): 120 cesium methanesulfonate, 10 CsCl, 10 HEPES, 10 EGTA, 5 Creatine Phosphate, 4 ATP-Mg, 0.4 GTP-Na (adjusted to pH 7.3-7.4 with CsOH).

Optically evoked synaptic currents were measured by delivering 2 light pulses (1 ms, spaced 50ms, generated by a CoolLED *p*E300) focused on the sample through the 40X immersion objective. Sweeps were delivered every 10 s. Synaptic responses were monitored while holding the membrane voltage at −70mV (for oEPSCs) and +10mV (for oIPSCs). We monitored different light stimulation intensities prior to baseline recording to achieve oEPSCs responses at ∼50% of the maximal response. Access resistance (10–30 MΩ) was monitored throughout the experiment, and recordings were discarded from further analysis if resistance changed >20%. We calculated the paired-pulse ratio (PPR) as the amplitude of the second peak divided by the amplitude of the first peak elicited by the twin stimuli. The excitation-inhibition (E/I) ratio was calculated by dividing the amplitude of the oEPSC at −70mV by the amplitude of the oIPSC at +10mV, during identical light intensity stimulation. To validate inhibitory and excitatory post-synaptic currents (PSCs) as gamma-aminobutyric acid (GABA)ergic and glutamatergic, respectively, in a subset of cells the GABAa receptor antagonist SR 95531 hydrobromide (gabazine, µM) was added to the bath while holding the cell at +10mV, or the AMPA receptor antagonist 6,7- Dinitroquinoxaline-2,3-dione (DNQX, µM) while holding the cell at −70mV. In another subset of cells, once the baseline measures were established, we tested for monosynaptic connectivity by bath application of 1µM Tetrodotoxin (TTX), followed by 100µM 4-Aminopyridine (4AP), and measured the amplitude of PSCs returning following 4-AP application. Based on the signal-to-noise ratio (S/N) of the recordings, currents under 5pA were considered unreliable and not considered further, as were currents rescued by 4AP application with an amplitude <10pA (non-monosynaptic; 2 instances, 1 HVC_X_ →HVC_X_, 1 HVC_X_→HVC_RA_).

### Histology and Immunohistochemistry

Birds were anesthetized with Euthasol (Virbac, TX, USA) and transcardially perfused with 4% paraformaldehyde in phosphate buffered saline (PBS). Free-floating sagittal sections (30 µm) were cut using a cryostat (Leica CM1950, Leica). Sections were first washed in PBS, then blocked in 3% bovine serum albumine in PBST (0.3% Triton X-100 in PBS) for 1 hour at RT and incubated with primary antibodies (α-NeuN MAB377 Millipore, α-GFP a11122 Invitrogen) diluted in the blocking buffer 4°C for 24 hours. Slices were then washed with PBS and incubated with fluorescent secondary antibodies (alexa-conjugated, Invitrogen, diluted in blocking buffer) at RT for 2 hours. After PBS wash, sections were mounted onto slides with Fluoromount-G (eBioscience, CA, USA). Composite images were acquired and stitched using an LSM 880 or LSM710 laser scanning confocal microscope (Carl Zeiss, Germany), and/or an Zeiss Axioscan Z1 (Univ of Texas Southwestern Medical Center Whole Brain Microscopy Facility, RRID:SCR_017949). Image analyses were performed using ImageJ. After electrophysiological recordings, the slices were incubated in 4% PFA in PBS. Sections were then washed in PBS, mounted on glass slides with Fluoromount-G (eBioscience, CA, USA), and visualized under an LSM 880 laser-scanning confocal microscope (Carl Zeiss, Germany).

### 3D brain imaging and processing

Imaging and processing of the sample brain with tracers injected in HVC (Alexa488-conjugated dextran 10000) and RA (Alexa568-conjugated dextran 10000) for 3D rendering were conducted with the help of Denise Ramirez and Ariana Nawaby (Univ of Texas Southwestern Medical Center Whole Brain Microscopy Facility, RRID:SCR_017949). After perfusion with 4%PFA the brain was embedded in oxidized agarose in preparation for sectioning. The TissueCyte 1000 instrument (TissueVision, Inc.) automatically sectioned the entire volume of the brain at 100mm in the coronal plane and collected mosaic image tiles encompassing each section.

Preprocessing: Images were downsampled to 1.5 um xy resolution and color contrast adjusted to provide high visual contrast between signals of interest and background.

Segmentation: A selected portion of signals of interest in the downsampled contrast adjusted images of the tissue were visually identified and annotated, then used to train a random forest classifier for segmentation in Ilastik (version 1.3.3)^59–62^. This classifier was applied to all section images in the brain to assign a probability score to each pixel in the image, corresponding to its chance of belonging to specific fluorescent signals, autofluoresence, or background noise.

The total autofluorescence, Alexa488 (green), and Alexa568 (red) pixelwise probability scores were further processed, then used for visualization.

Segmentation Post-Processing: In order to create a grey silhouette of the overall shape of the brain, the autofluoresence probability signal was thresholded using the ImageJ default thresholding algorithm. Any holes in the binary mask were then flood filled, and particles greater than 3024 px^2 were removed. Green and red probabilities were thresholded at 105 8bit pixel intensity, and 79 8bit pixel intensity, respectively, determined visually to reduce low probability noise in the image. GFP signal in the rostral most portion of the brain (past section 135) was dimmed for better visibility of more caudal structures, by subtracting the pixel intensities by 140 pixel intensity units in the 8-bit range.

Visualization: Combined RGB images of the autofluoresence (grey), Alexa488 (green), and Alexa568 (red) post-processed probabilities were visualized in 3D using VAA3D software (version: V3.447, https://home.penglab.com/proj/vaa3d/home/index.html).

### Song analysis

Bird songs were recorded and analyzed using Sound Analysis Pro 2011 (SAP, Tchernichovski et al., 2000) and plots were made with a modified version of Avian Vocalization Network^63^ (AVN). We manually measured and categorized the outcomes of optogenetic stimulations. Truncations were defined as stimulation-contingent (≤300ms) atypical amplitude decays (not present in control motifs) visible as silent gaps in the spectrogram. Truncation latencies were measured from the onset of the light delivery to the onset of the optically-contingent silent gap. Stop is defined as truncation not followed by continuation/resumption of the motif. Twenty stimulated song segments were measured for stimulated and non-stimulated conditions for quantification of acoustic properties and sound similarity (SAP). Acoustic properties of the stimulated segment were measured and compared to the corresponding song fragment in unstimulated control motifs. When optical stimulation did not cause truncation, acoustic properties were calculated on the song fragment from the onset of optical stimulation to the end of the last syllable. The entire motif was analyzed during 1s stimulation trials.

In the 1s time window after song truncation, optical stimulation effects were manually classified as falling into one of four categories:1) motif reset (i.e., re-starting with the first song syllable, with introductory notes or with syllables that normally link motifs), 2) calls (typical zebra finch calls), 3) introductory notes (introductory notes not followed by motif initiation), or 4) pause and continuation (post-truncation motif resumption at any syllable in the motif other than the first syllable). To calculate the normalized motif reset probability, the number of motifs/bout was calculated over 30-50 bouts (defined as chains of motifs, started with introductory notes and mostly uninterrupted; in rare occasions we found motifs produced within 1s from other motifs, and they were considered as part of the previous bout. M=average motifs-per-bout). Each bird’s probability of motif truncation was then normalized (normalized motif reset probability = motif reset probability/[1-(1/M] following the logic that 1/M is the likelihood of each motif to be the last in the bout and not be followed by another motif, and therefore 1-(1/M) is the probability of a motif to be followed by another motif in the current bout. The probability of reset implies the presence of a motif after the truncated one examined, therefore dividing by the likelihood of that motif being followed by another one returns a normalized measure of the reset).

To report cross-motif quantification of truncation or reset latency and resumed vocalization identity probability, events were categorized depending on the timepoint within the motif at which the onset of the corresponding stimulation occurred. The events were then grouped in 10% bins across the motif duration, per each bird, to allow for comparison between birds with different motif lengths. 100% for each bird was set to the duration of the motif −100ms, as the latency to truncation when applied later than 100ms before the end of the motif would lead to unclear effects on the syllables (average truncation latency across groups=74.36±3.06). Whenever the stimulation happened in the last 100ms of motif the events were classified in the −20 - 0% bins, affecting the transition to the following motif (if any). Stimulations, truncations and post-truncation effects occurring during intro notes and inter-motif connecting syllables were assigned to these −20 – 0% time bins based on their temporal distance to the syllable A (if no syllable A onset was produced the effects were not considered for further analysis, as we couldn’t categorize the intro note as produced at specific distance from the motif for the % computation).

To evaluate the likelihood of optogenetic inhibition or stimulation across a motif-motif transition to terminate a bout (Fig. 3K and Extended Data Fig. 3D) we delivered light or sham stimulation across the motif and extending after its end, and quantified the probability of the stimulation to be contingent with the termination of the bout for 50 trials in each condition.

In lesion experiments, a minimum of 50 motifs were scored with SAP against pre-surgery motifs. Failed motif starts were defined as a series of introductory notes not leading to a motif. The number of motifs in a bout were counted over 50 bouts. In case of absence of motifs being produced (the bird did not sing) the accuracy was assigned the value of 0 for sake of classification.

### Recurrent Circuit Model of HVC

The computational model employed in this study is based on a canonical recurrent circuit model, the continuous attractor neural network (CAN). In a typical CAN, excitatory neurons are arranged to uniformly cover a linear feature space (e.g., the location of the timing chain in the current example, and have mutual interactions via recurrent connections). This configuration gives rise to a continuous manifold that sustains a series of activity bumps. A song motif is considered to be controlled by an activity bump traversing from one end of the chain to another.

To better reflect the biological characteristics of the songbird HVC, we introduced several specific features. The model incorporates five distinct neuron types to capture the functional diversity in the songbird HVC.

1. Excitatory neurons (HVC_RA_, 𝐫_RA_, and HVC_X_, 𝐫_X_)

The excitatory neurons responsible for encoding the neural sequence are divided into two groups: HVC_RA_ and HVC_X_, with their firing rates denoted as 𝐫_RA_and 𝐫_X_, respectively. Consistent with experimental observations, the model only includes inter-group connections, and leaves neurons within the same group unconnected. Simulations demonstrated that these inter-group connections are sufficient to self-sustain non-zero responses and moving sequences.

1. Global inhibitory neurons (𝐫*_g_*)

To keep the stability of the network, the network model contains a global inhibitory neuron with the firing rate 𝐫_g_. Compared with excitatory neurons, in the model this neuron has more rapid dynamics and a steeper activation function to provide effective global inhibition.

1. Local inhibitory neurons (𝐫_I_)

The circuit model has another group of inhibitory neurons (𝐫_I_) providing local, structured inhibitory feedback to the excitatory populations, which is essential to generate spontaneous movement of excitatory neurons’ population activity bumps within the circuit. The 𝐫_I_ bump slightly lags behind the excitatory neuron bumps due to transmission delay and slow dynamics, so that the E neurons at more distant locations will be suppressed less and build up more activities. As a result, the E neurons’ activity bump is ‘pushed’ to move forward.

1. Peri-song neurons (𝐫_ps_)

The circuit model also contains an HVC_RA_ peri-song neuron group^44^ (𝐫_ps_) that is modeled to target HVC_RA_ song neurons at the initial end of the manifold. This group plays a critical role in initiating and resetting motif generation.

### Circuit dynamics

The neural dynamics underlying these activities are captured by a set of dynamic equations:

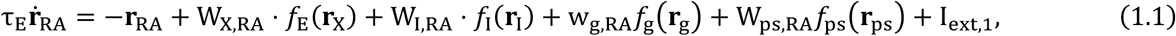

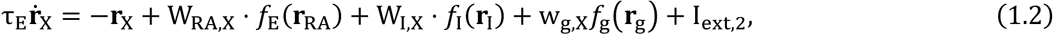

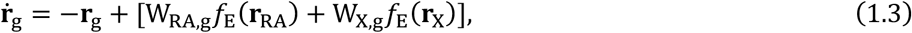

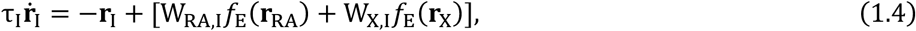

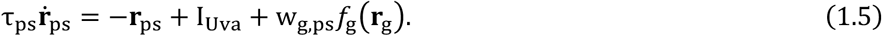

In these equations, subscripts denote the neuron types. The parameter 𝜏 represents the time constant and 𝑓(⋅) denotes the activation function for each neuron group. External input currents are denoted as I_ext_, and specific terms such as I_Uva_ correspond to input from Uva. The capital W_A,B_ indicates the connection matrix from group A to B with dimensions N_B_ × N_A_, where N is the number of neurons in the respective group, while the lowercase w indicates the scalar connection strength. For convenience, we set N_RA_ = N_X_ = N_I_ = N and N_g_ = N_ps_ = 1. Specifically, to support a continuous manifold, the entries of connections between excitatory and local inhibitory neurons are determined by the distance between the index of pre- and post-synaptic neurons

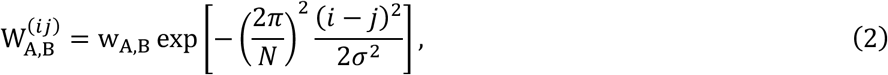

where w_AB_ (A, B ∈ Error! Bookmark not deVined.) denotes the peak weight of the weight from neuronal population A to B.

To target the peri-song output to the initial end of the manifold, W_ps,E_ is a N × 1 matrix with its column in a Gaussian profile centering at 0:

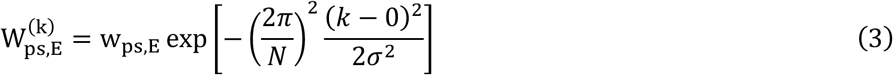

### Sequence initiation

The fundamental property of the network is its ability to spontaneously generate neural sequences. In our model, peri-song neurons initiate the sequential activity. The peri-song neurons receive excitatory input, likely originating from the upstream nucleus Uva, while simultaneously receiving inhibitory input from the global inhibitory neurons.

When the network is silenced, whether at rest or following truncation, activity in the global inhibitory neuron decreases, which disinhibits the peri-song neurons. This release from inhibition then triggers the onset of a motif.

### Boundaries

Following the activation of excitatory neurons, the activity bump is driven by locally structured inhibitory feedback from 𝐫_&_ to traverse the continuous manifold. For the bump to gain a directional motion tendency, the inhibitory feedback is intentionally enhanced at the initial locations on the chain. Due to the recurrent nature of the network, the bump would ordinarily “bounce” back upon reaching the end of the chain; however, this behavior is inconsistent with observed data. To address this, we introduced a fading mechanism for excitatory-to-excitatory (E-E) connections as the bump approaches the boundary, simulating a “boundary effect.” This gradual reduction in connectivity causes the bump to diminish as it reaches the endpoint, resulting in an automatic cessation of activity that mimics the natural termination of a motif. These two boundary behaviors were implemented by multiplying the connection strength with a compensation factor:

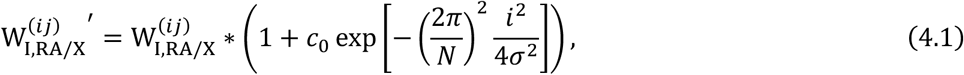

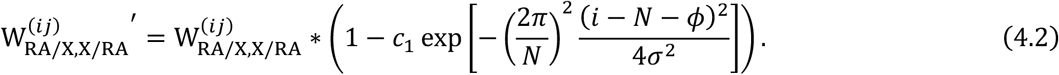

𝜙 is an offset term, in which we take the value 𝜙 = 0.5𝜎𝑁/2𝜋. The compensated connection matrices are shown in Extended Data Fig 9C,D.

### Truncation

To simulate optogenetic stimulation truncating HVC neuronal sequences observed in experimental studies, we applied an intense, spatially homogeneous pulse input to eitherHVC_RA_or HVC_X_ neurons. Following this stimulation, both 𝐫_RA_ and 𝐫_X_ became hyper-activated, leading to rapid suppression by the fast response of 𝐫_g_. These neurons remain suppressed until 𝐫_g_ activity subsides, corresponding to the observed motif truncation (Fig.7, Supp Fig. 2). Subsequently, the peri-song neurons re-initiate the neural sequence, allowing the motif to resume from the beginning. Considering that in the current model HVC_RA_ and HVC_X_ are connected symmetrically, we only simulated optogenetic stimulation on HVC_RA_ as a verification.

### HVC_X_ degradation

To replicate the effects of TeNT expression in HVC_X_ neurons, as observed in Fig. 6B, we manually modified the output projections of HVC_X_ neurons to simulate their degradation. Let 𝑝 denote the proportion of degradation. Under this condition, the degraded projection from HVC_X_ to HVC_RA_ (W_X,RA′_) can be expressed as:

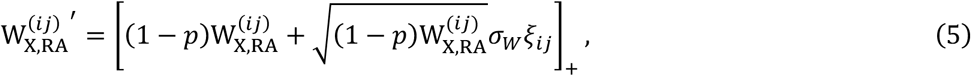

where 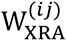 represents the original connection strength, 𝜎*_W_* denotes the variation coefficient, and 𝜉*_ij_* is an independent Gaussian noise term indexed by pre- and post- neuron index ij. [*x*]_*_ = max(*x*, 0) denotes the negative rectification, ensuring the weight is always excitatory (positive).

During the synaptic degradation over weeks, experiments found the neuronal sequences in different trials observed within the same day could transverse and then disappear at different random locations and on the chain. We assume the synaptic weights during the same day is nearly the same, and then the random progression along the chain is resulted from the variability of single neurons. Therefore, to reproduce the random progression along the chain during synaptic degradation, each HVC_RA_ neuron **r**_RA_(*j*) receives a Poisson-like noise *I_noise_*, mimicking stochastic spike generation,

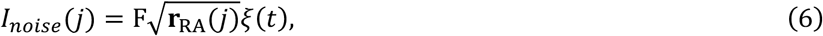

where F is the fano factor scaling the noise, and 𝜉(*t*) is a standard Gaussian white noise. Moreover, the noises received by different neurons are independent with each other. Under these conditions, we observed that sequences terminated at random positions. As illustrated in Fig. 7E, the average sequence length decreased as the proportion of neuronal degradation increased.

## Statistical Analysis

All data were analyzed with GraphPad Prism. Data were tested for normality using the Shapiro-Wilk Test. Parametric and non-parametric statistical tests were used. To compare between 2 groups, *t-test*, *Mann-Whitney* and *Kolmogorov-Smirnov* tests were used. For more than two conditions, *one*- and *two-way ANOVAs* or Kruskal-Wallis test were performed. Cumulative probability curves were calculated for each animal, then tested in groups for statistical significance. Only one comparison among all groups was made, to avoid repeatedly comparing the same dataset (i.e. HVC) to single other data sets. *Fisher* or *X^2^* tests, followed by *Dunn’s* post-hoc test, were used to compare the probability of finding optically evoked responses across the HVCPN classes while stimulating the different afferents. *Dunn’s*, *Sidak’s* or *Holm-Sidak’s* post-hoc tests were used to correct for multiple comparisons Statistical significance refers to \**P* < 0.05, ** *P* < 0.01, *** *P* < 0.001.

## Author Contributions

MT and TFR conceived the project. MT, JZ, DHA, ESM, TMIK, JC, HP, BGC, and WZ designed the methodology. MT, JZ, BGC, and WZ performed the investigation. MT, JZ, ESM, TMIK, and BGC visualized the project. TFR aquired funding. TFR administered and supervised the project. MT, WZ, BGC, and TF wrote the original draft of the manuscript. All authors contributed to writing, reviewing and editing the manuscript. Authors declare no competing interests. All data is available in the main text or the supplementary materials.

## Acknowledgments

We thank Drs. Timothy Gentner, William Dauer, Daisuke Hattori, Sebastian Choi, and members of the Roberts lab for comments on an initial version of the manuscript and Drs. Michael Long, Marc Schmidt, and Dimitri Aronov for valuable discussions of our data as the project unfolded. We thank Jennifer Holdway, Luis Garcia, and Rico Cabuco for laboratory support. We thank Dr. Denise Ramirez and Ariana Nawaby (Univ of Texas Southwestern Medical Center Whole Brain Microscopy Facility, RRID:SCR_017949) for the 3D imaging and rendering. We thank Jesse Hilton, Richonda Hunte, and Pamela Jennings for administrative support. We thank the National Institutes of Health for supporting this research through grants UF1NS115821 and R01NS108424 to TFR and F99NS124172 to DHA.

## References

1 Doupe, A. J. & Kuhl, P. K. Birdsong and human speech: common themes and mechanisms. Annu Rev Neurosci 22, 567–631 (1999).

2 Konopka, G. & Roberts, T. F. Insights into the Neural and Genetic Basis of Vocal Communication. Cell 164, 1269–1276 (2016). 10.1016/j.cell.2016.02.039

3 Brainard, M. S. & Doupe, A. J. What songbirds teach us about learning. Nature 417, 351–358 (2002).

4 Hahnloser, R. H., Kozhevnikov, A. A. & Fee, M. S. An ultra-sparse code underlies the generation of neural sequences in a songbird. Nature 419, 65–70 (2002). 10.1038/nature00974

5 Fee, M. S., Kozhevnikov, A. A. & Hahnloser, R. H. Neural mechanisms of vocal sequence generation in the songbird. Ann N Y Acad Sci 1016, 153–170 (2004).

6 Long, M. A., Jin, D. Z. & Fee, M. S. Support for a synaptic chain model of neuronal sequence generation. Nature 468, 394–399 (2010). 10.1038/nature09514

7 Lynch, G. F., Okubo, T. S., Hanuschkin, A., Hahnloser, R. H. & Fee, M. S. Rhythmic Continuous-Time Coding in the Songbird Analog of Vocal Motor Cortex. Neuron 90, 877–892 (2016). 10.1016/j.neuron.2016.04.021

8 Picardo, M. A. et al. Population-Level Representation of a Temporal Sequence Underlying Song Production in the Zebra Finch. Neuron 90, 866–876 (2016). 10.1016/j.neuron.2016.02.016

9 Ashmore, R. C., Renk, J. A. & Schmidt, M. F. Bottom-up activation of the vocal motor forebrain by the respiratory brainstem. J Neurosci 28, 2613–2623 (2008).10.1523/JNEUROSCI.4547-07.2008

10 Ashmore, R. C., Wild, J. M. & Schmidt, M. F. Brainstem and forebrain contributions to the generation of learned motor behaviors for song. J Neurosci 25, 8543–8554 (2005).

11 Hamaguchi, K., Tanaka, M. & Mooney, R. A Distributed Recurrent Network Contributes to Temporally Precise Vocalizations. Neuron 91, 680–693 (2016). 10.1016/j.neuron.2016.06.019

12 Moll, F. W. et al. Thalamus drives vocal onsets in the zebra finch courtship song. Nature 616, 132–136 (2023). 10.1038/s41586-023-05818-x

13 Long, M. A. & Fee, M. S. Using temperature to analyse temporal dynamics in the songbird motor pathway. Nature 456, 189–194 (2008).

14 Elmaleh, M., Kranz, D., Asensio, A. C., Moll, F. W. & Long, M. A. Sleep replay reveals premotor circuit structure for a skilled behavior. Neuron 109, 3851–3861.e3854 (2021). 10.1016/j.neuron.2021.09.021

15 Armstrong, E. & Abarbanel, H. D. Model of the songbird nucleus HVC as a network of central pattern generators. J Neurophysiol 116, 2405–2419 (2016). 10.1152/jn.00438.2016

16 Schmidt, M. F. Pattern of interhemispheric synchronization in HVc during singing correlates with key transitions in the song pattern. J Neurophysiol 90, 3931–3949 (2003). 10.1152/jn.00003.2003

17 Amador, A., Perl, Y. S., Mindlin, G. B. & Margoliash, D. Elemental gesture dynamics are encoded by song premotor cortical neurons. Nature 495, 59–64 (2013). 10.1038/nature11967

18 Danish, H. H., Aronov, D. & Fee, M. S. Rhythmic syllable-related activity in a songbird motor thalamic nucleus necessary for learned vocalizations. PloS one 12, e0169568 (2017). 10.1371/journal.pone.0169568

19 Vu, E. T., Mazurek, M. E. & Kuo, Y. C. Identification of a forebrain motor programming network for the learned song of zebra finches. J Neurosci 14, 6924–6934 (1994). 10.1523/jneurosci.14-11-06924.1994

20 Vu, E. T., Schmidt, M. F. & Mazurek, M. E. Interhemispheric coordination of premotor neural activity during singing in adult zebra finches. J Neurosci 18, 9088–9098 (1998). 10.1523/jneurosci.18-21-09088.1998

21 Ashmore, R. C., Bourjaily, M. & Schmidt, M. F. Hemispheric coordination is necessary for song production in adult birds: implications for a dual role for forebrain nuclei in vocal motor control. J Neurophysiol 99, 373–385 (2008).10.1152/jn.00830.2007

22 Arfin, S. K., Long, M. A., Fee, M. S. & Sarpeshkar, R. Wireless neural stimulation in freely behaving small animals. J Neurophysiol 102, 598–605 (2009). 10.1152/jn.00017.2009

23 Histed, M. H., Bonin, V. & Reid, R. C. Direct activation of sparse, distributed populations of cortical neurons by electrical microstimulation. Neuron 63, 508–522 (2009). 10.1016/j.neuron.2009.07.016

24 Marshel, J. H. et al. Cortical layer-specific critical dynamics triggering perception. Science 365 (2019). 10.1126/science.aaw5202

25 Hamaguchi, K., Tschida, K. A., Yoon, I., Donald, B. R. & Mooney, R. Auditory synapses to song premotor neurons are gated og during vocalization in zebra finches. eLife 3, e01833 (2014).10.7554/eLife.01833

26 Vallentin, D. & Long, M. A. Motor origin of precise synaptic inputs onto forebrain neurons driving a skilled behavior. J Neurosci 35, 299–307 (2015). 10.1523/JNEUROSCI.3698-14.2015

27 Kozhevnikov, A. A. & Fee, M. S. Singing-related activity of identified HVC neurons in the zebra finch. J Neurophysiol 97, 4271–4283 (2007).

28 Marder, E. & Bucher, D. Central pattern generators and the control of rhythmic movements. Curr Biol 11, R986–996 (2001). 10.1016/s0960-9822(01)00581-4

29 Berkowitz, A. Expanding our horizons: central pattern generation in the context of complex activity sequences. J Exp Biol 222 (2019). 10.1242/jeb.192054

30 Coleman, M. J. & Vu, E. T. Recovery of impaired songs following unilateral but not bilateral lesions of nucleus uvaeformis of adult zebra finches. J Neurobiol 63, 70–89 (2005).

31 Trusel, M. et al. Synaptic Connectivity of Sensorimotor Circuits for Vocal Imitation in the Songbird. bioRxiv, 2022.2011.2008.515692 (2024). 10.1101/2022.11.08.515692

32 Coleman, M. J., Roy, A., Wild, J. M. & Mooney, R. Thalamic gating of auditory responses in telencephalic song control nuclei. J Neurosci 27, 10024–10036 (2007). 10.1523/jneurosci.2215-07.2007

33 Cynx, J. Experimental determination of a unit of song production in the zebra finch (Taeniopygia guttata). J Comp Psychol 104, 3–10 (1990). 10.1037/0735-7036.104.1.3

34 Franz, M. & Goller, F. Respiratory units of motor production and song imitation in the zebra finch. J Neurobiol 51, 129–141 (2002).

35 Williams, H. & Vicario, D. S. Temporal patterning of song production: participation of nucleus uvaeformis of the thalamus. J Neurobiol 24, 903–912 (1993). 10.1002/neu.480240704

36 Aronov, D., Andalman, A. S. & Fee, M. S. A specialized forebrain circuit for vocal babbling in the juvenile songbird. Science 320, 630–634 (2008).10.1126/science.1155140

37 Fetterman, G. C. & Margoliash, D. Rhythmically bursting songbird vocomotor neurons are organized into multiple sequences, suggesting a network/intrinsic properties model encoding song and error, not time. bioRxiv (2023).10.1101/2023.01.23.525213

38 Fee, M. S. & Goldberg, J. H. A hypothesis for basal ganglia-dependent reinforcement learning in the songbird. Neuroscience 198, 152–170 (2011). 10.1016/j.neuroscience.2011.09.069

39 Scharg, C., Kirn, J. R., Grossman, M., Macklis, J. D. & Nottebohm, F. Targeted neuronal death agects neuronal replacement and vocal behavior in adult songbirds. Neuron 25, 481–492 (2000).

40 Sanchez-Valpuesta, M. et al. Corticobasal ganglia projecting neurons are required for juvenile vocal learning but not for adult vocal plasticity in songbirds. Proc Natl Acad Sci U S A 116, 22833–22843 (2019). 10.1073/pnas.1913575116

41 Mooney, R. & Prather, J. F. The HVC microcircuit: the synaptic basis for interactions between song motor and vocal plasticity pathways. J Neurosci 25, 1952–1964 (2005). 10.1523/jneurosci.3726-04.2005

42 Kosche, G., Vallentin, D. & Long, M. A. Interplay of inhibition and excitation shapes a premotor neural sequence. J Neurosci 35, 1217–1227 (2015). 10.1523/jneurosci.4346-14.2015

43 Petreanu, L., Mao, T., Sternson, S. M. & Svoboda, K. The subcellular organization of neocortical excitatory connections. Nature 457, 1142–1145 (2009). 10.1038/nature07709

44 Daliparthi, V. K. et al. Transitioning between preparatory and precisely sequenced neuronal activity in production of a skilled behavior. eLife 8 (2019). 10.7554/eLife.43732

45 Sadeh, S. & Clopath, C. Inhibitory stabilization and cortical computation. Nat Rev Neurosci 22, 21–37 (2021). 10.1038/s41583-020-00390-z

46 Bera, K., Shukla, A. & Bapi, R. S. Motor Chunking in Internally Guided Sequencing. Brain Sci 11 (2021). 10.3390/brainsci11030292

47 Tosatto, L., Fagot, J., Nemeth, D. & Rey, A. Chunking as a function of sequence length. Anim Cogn 28, 2 (2024). 10.1007/s10071-024-01835-z

48 Fonollosa, J., Neftci, E. & Rabinovich, M. Learning of Chunking Sequences in Cognition and Behavior. PLoS Comput Biol 11, e1004592 (2015). 10.1371/journal.pcbi.1004592

49 Okubo, T. S., Mackevicius, E. L., Payne, H. L., Lynch, G. F. & Fee, M. S. Growth and splitting of neural sequences in songbird vocal development. Nature 528, 352–357 (2015). 10.1038/nature15741

50 Piristine, H. C., Choetso, T. & Gobes, S. M. A sensorimotor area in the songbird brain is required for production of vocalizations in the song learning period of development. Dev Neurobiol 76, 1213–1225 (2016). 10.1002/dneu.22384

51 Foster, E. F. & Bottjer, S. W. Lesions of a telencephalic nucleus in male zebra finches: Influences on vocal behavior in juveniles and adults. J Neurobiol 46, 142–165 (2001). 10.1002/1097-4695(20010205)46:2<142::aid-neu60>3.0.co;2-r

52 Roberts, T. F., Gobes, S. M., Murugan, M., Olveczky, B. P. & Mooney, R. Motor circuits are required to encode a sensory model for imitative learning. Nat Neurosci 15, 1454–1459 (2012). 10.1038/nn.3206

53 Roberts, T. F. et al. Identification of a motor-to-auditory pathway important for vocal learning. Nat Neurosci 20, 978–986 (2017). 10.1038/nn.4563

54 Roberts, T. F. & Mooney, R. Motor circuits help encode auditory memories of vocal models used to guide vocal learning. Hear Res 303, 48–57 (2013). 10.1016/j.heares.2013.01.009

55 Roberts, T. F., Klein, M. E., Kubke, M. F., Wild, J. M. & Mooney, R. Telencephalic neurons monosynaptically link brainstem and forebrain premotor networks necessary for song. J Neurosci 28, 3479–3489 (2008).

56 Stauger, T. R. et al. Axial organization of a brain region that sequences a learned pattern of behavior. J Neurosci 32, 9312–9322 (2012). 10.1523/JNEUROSCI.0978-12.2012

57 Xiao, L. et al. A Basal Ganglia Circuit Sugicient to Guide Birdsong Learning. Neuron 98, 208–221.e205 (2018). 10.1016/j.neuron.2018.02.020

58 Ali, F. et al. The basal ganglia is necessary for learning spectral, but not temporal, features of birdsong. Neuron 80, 494–506 (2013). 10.1016/j.neuron.2013.07.049

59 Peng, H., Bria, A., Zhou, Z., Iannello, G. & Long, F. Extensible visualization and analysis for multidimensional images using Vaa3D. Nature Protocols 9, 193–208 (2014). 10.1038/nprot.2014.011

60 Peng, H., Ruan, Z., Long, F., Simpson, J. H. & Myers, E. W. V3D enables real-time 3D visualization and quantitative analysis of large-scale biological image data sets. Nature Biotechnology 28, 348–353 (2010). 10.1038/nbt.1612

61 Peng, H. et al. Virtual finger boosts three-dimensional imaging and microsurgery as well as terabyte volume image visualization and analysis. Nature Communications 5, 4342 (2014). 10.1038/ncomms5342

62 Berg, S. et al. ilastik: interactive machine learning for (bio)image analysis. Nature Methods 16, 1226–1232 (2019). 10.1038/s41592-019-0582-9

63 Koch, T. M. I., Marks, E. & Roberts, T. F. AVN: A Deep Learning Approach for the Analysis of Birdsong. bioRxiv, 2024.2005.2010.593561 (2024). 10.1101/2024.05.10.593561

